# Molecular mechanisms underlying amyloid lowering by aducanumab: differential and comparative effects of sex and IgG reveal the post-treatment disease brain

**DOI:** 10.64898/2026.06.16.731973

**Authors:** KA Haynes, RS Pandey, EH Doud, ZA Cope, GJ Little, S-P Williams, U Nepali, SK Quinney, A Nagar, NB Charbe, L da Silva, JL Dage, DM Duong, NT Seyfried, M Sasner, BT Lamb, AL Oblak, PR Territo, GW Carter, SJ Sukoff Rizzo

## Abstract

**INTRODUCTION:** Improving the predictive validity of preclinical studies for Alzheimer’s disease (AD) requires rigorous evaluation of therapeutic efficacy, safety, and sex-specific responses in translationally relevant models. As amyloid-targeting monoclonal antibodies continue to advance clinically, there is an urgent need to define the molecular milieu that persists after amyloid is reduced and disease progression continues. Leveraging the NIA-funded MODEL-AD Preclinical Testing Core, we investigated the biochemical, functional, and multi-omic signatures associated with chronic administration of murine chimeric aducanumab (chAdu) in 5XFAD mice, including the contribution of IgG-mediated effects.

**METHODS:** Male and female 5XFAD mice were treated weekly with chAdu beginning at 8 months of age and compared to age-and sex-matched murine IgG2aκ isotype (IgG) and saline controls. Plasma and brain pharmacokinetics, amyloid-beta (Aβ), behavioral assessments, and treatment-emergent anti-drug antibodies (ADAs) were quantified. Post-treatment transcriptomic and proteomic analyses were performed to assess molecular pathways associated with chAdu and IgG exposure following 17-week treatment.

**RESULTS:** chAdu produced sex-dependent changes in Aβ, including increased plasma Aβ42:40 and reductions in brain Aβ which were associated with mild behavioral impairments in the absence of improvements in cognitive function. IgG control treatment produced similar reductions, indicating biologically active IgG-mediated processes independent of Aβ-targeted specificity. Treatment-emergent ADAs occurred in 10% of chAdu-treated mice and were associated with reduced drug exposure and efficacy. Multi-omics analyses confirmed sex-dependent and IgG-mediated effects at both the transcriptomic and proteome level revealing disease-associated genes and proteins not altered despite reductions in amyloid with treatment.

**DISCUSSION:** These findings demonstrate sex-dependent PK and pharmacodynamic responses to chAdu, identify biologically meaningful IgG-driven effects, and reveal molecular signatures that persist after amyloid reduction. This work provides biological insights into pathways that may remain insufficiently addressed following amyloid lowering; revealing novel targets for future drug discovery to prevent and treat disease.

## INTRODUCTION

Despite substantial advances in understanding the pathophysiology of Alzheimer’s disease (AD), most therapeutic approaches have failed to meaningfully slow cognitive decline or halt disease progression [1,2].Amyloid beta (Aβ)-targeting monoclonal antibodies (mAbs) were the first disease-modifying therapies approved for AD [3,4]. Aduhelm™ (aducanumab) received accelerated approval from the U.S. Food and Drug Administration in 2021 based on amyloid plaque reduction despite significant rates of amyloid-related imaging abnormalities [5–7]. Although Aduhelm™ was later withdrawn from the market, subsequent approvals of lecanemab (Leqembi®) and donanemab (Kisunla™) reflect continued clinical reliance on amyloid-lowering immunotherapies [8–10]. While these therapies effectively reduce amyloid plaques and exhibit improved safety profiles, they do not prevent continued clinical decline [10–13]. This highlights a major unmet need to understand what comprises the diseased brain milieu when amyloid aggregates are minimized yet the disease continues to progress.

To address this knowledge gap, we characterized biochemical, behavioral, pharmacokinetic, and molecular responses to chronic Aβ-targeting antibody exposure in aged 5XFAD mice with established amyloid pathology using the drug-screening resources of the MODEL-AD Preclinical Testing Core (PTC) [1,14–16]. We chronically administered murine chimeric aducanumab (chAdu), representing the class of Aβ-targeting IgG antibodies, and compared its effects with IgG isotype control treatment in amyloidogenic 5XFAD mice [15]. We evaluated the effects of chAdu and IgG on plasma and brain Aβ, behavior, treatment-emergent anti-drug antibodies (ADAs), and post-treatment proteomic and transcriptomic signatures to characterize molecular alterations that persist following amyloid lowering and identify candidate pathways for future therapeutic interventions.

## METHODS

All raw data and methodological details are available via the AD Knowledge Portal: https://doi.org/10.7303/syn74927054. Additional detailed methods are provided in the Supplementary Methods.

### Subjects

All experiments were reviewed and approved by the Institutional Animal Care and Use Committee (IACUC) at the University of Pittsburgh. Male hemizygous 5XFAD mice (JAX stock #34848) were crossed with female C57BL/6J mice (JAX stock #000664) to generate hemizygous 5XFAD and wildtype (WT) littermate controls, as previously described [15,16]. Subjects entered studies at 8-9 months of age, when substantial amyloid pathology is present [15].

In accordance with ARRIVE guidelines [17], treatment groups were randomized and counterbalanced prior to dosing, and genotype and treatment assignments remained blinded throughout treatment, sample processing, and data analysis. Housing conditions, genotyping procedures, exclusion criteria, and cohort samples sizes are provided in the Supplementary Methods and Tables S1-S3.

### Drug characterization and immunogenicity screening

Murine chimeric aducanumab (chAdu) was produced as previously described [18], and murine IgG2aκ (IgG) was purchased from Crown Bioscience (Product Code C0006). Plasma and brain chAdu concentrations were quantified by PRM mass spectrometry [18,19], and ADAs were assessed using ACE bridging ELISA [20,21].

### Experimental design

As illustrated in Figure 1A, the study design included a 4-week pilot pharmacokinetic (PK) study and a 17-week chronic treatment study. Chronic dose levels were guided by previously published efficacy studies of chimeric aducanumab in Tg2576 mice [6]. The pilot study characterized chAdu and informed population PK (popPK) modeling. The chronic study evaluated biochemical, behavioral, and multi-omic outcomes following 17-week treatment, with behavioral assessments initiated after 12 weeks.

**Figure 1.**
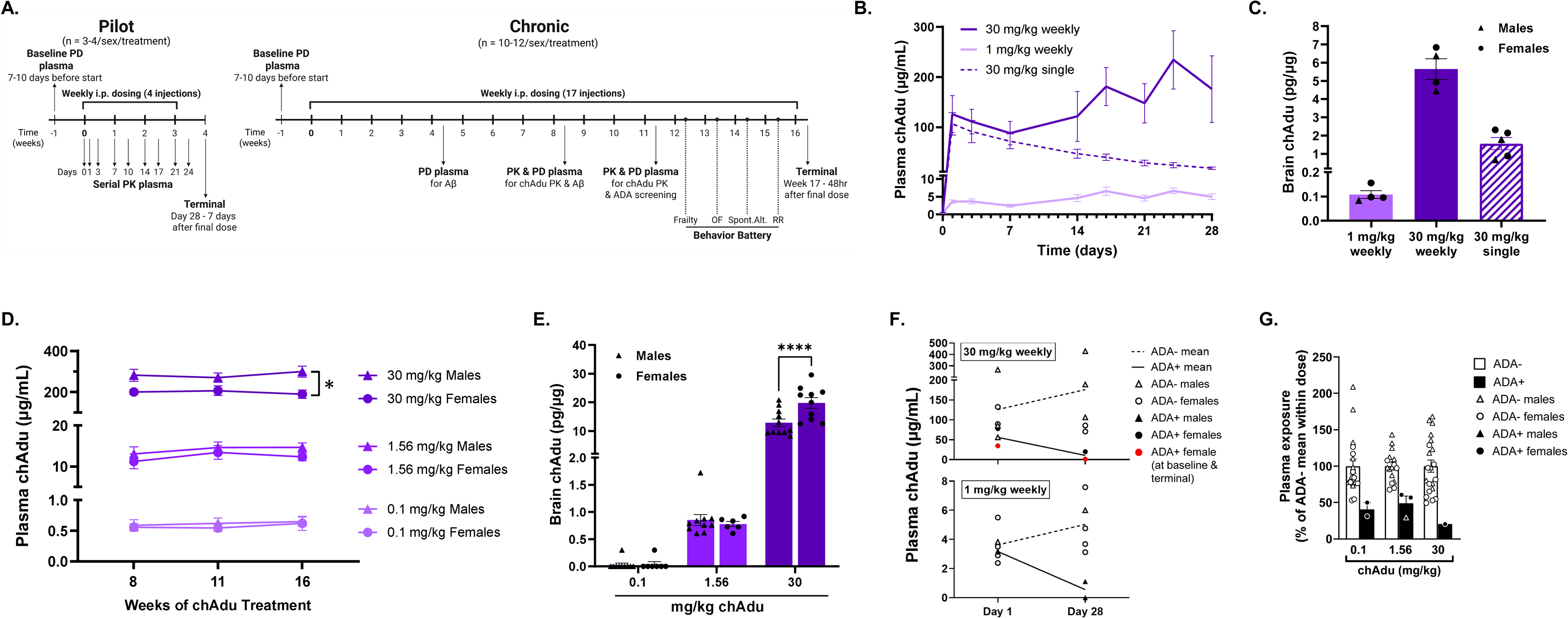
Pharmacokinetics and development of anti-drug antibodies (ADA) in response to aducanumab treatment for 4-weeks versus 17-weeks in aged 5XFAD mice. These studies evaluated response to treatment with murinized chimeric aducanumab (chAdu) administered once weekly, intraperitoneally (i.p.) to aged male and female 5XFAD mice. (A) Study design for 4-week pilot and 17-week chronic treatment cohorts, including dosing schedules and treatment groups, sample collection timepoints, and behavioral assessments in 5XFAD mice. Panel A was created with BioRender.com. (B) Plasma chAdu concentrations following 4-week treatment in 5XFAD mice. Plasma concentrations were measured longitudinally. Data represent mean ± SEM with sexes pooled. (C) ChAdu brain concentrations following 4-week treatment. Symbols represent individual mice (triangles, males; circles, females); bars indicate mean ± SEM. (D) Longitudinal plasma chAdu concentrations over 17-week treatment period. Within each dose group, data were analyzed by two-way repeated-measures ANOVA with Geisser–Greenhouse correction and Šídák’s multiple comparisons test. Symbols indicate sex (triangles, males; circles, females), and points indicate mean ± SEM. (\**p* < 0.05). (E) Brain chAdu concentrations following once weekly 17-week treatment in 5XFAD mice. Data were analyzed by two-way ANOVA with Šídák’s multiple comparisons test between sexes within each dose group. Symbols represent individual mice (triangles, males; circles, females); bars indicate mean ± SEM. (\*\*\*\**p* < 0.0001). (F) Longitudinal plasma chAdu concentrations following 4-week treatment stratified by terminal ADA status. Lines represent group means from repeated measurements within subjects. N=1 subject screened ADA positive (ADA+) prior to dosing (red symbols). Symbols represent individual mice (triangles, males; circles, females). (G) Plasma chAdu concentrations were calculated within each dose group and expressed as a percentage of the ADA-mean (set to 100%). ADA+ and ADA-groups were compared across all dose groups using a two-tailed Mann–Whitney test. Symbols represent individual animals (triangles, males; circles, females); bars indicate mean ± SEM.

### Chronic treatment of chAdu and IgG

Male and female 5XFAD mice received once-weekly intraperitoneal administration of chAdu (0.1, 1.56, or 30 mg/kg), IgG isotype control (1.56 mg/kg), or saline for 17 weeks (n = 10–12/sex/group; Fig. 1A). Age-and sex-matched WT littermates received IgG and served as behavioral controls.

The chronic study was conducted in two balanced cohorts staggered four weeks apart. Longitudinal plasma samples were collected for PK, pharmacodynamic, and immunogenicity analyses, and terminal plasma, CSF, and brain tissues were collected 48 hours after the final dose.

### Brain and plasma analysis of Aβ42 and Aβ40

Soluble and insoluble brain protein fractions were prepared as previously described [16,22]. Aβ40 and Aβ42 concentrations were quantified in brain and plasma using the Meso Scale Discovery V-PLEX Aβ Peptide Panel 1 (4G8) (#K15199E).

### Behavioral assessments

Behavioral assessments (frailty, open field, spontaneous alternation, and rotarod) were performed between treatment weeks 13 and 16 (Fig. 1A). Only ADA-negative (ADA-) animals were included in behavioral analyses.

### Statistics

Statistical analyses were performed using GraphPad Prism (v10) with ANOVA-based and nonparametric approaches as appropriate. Primary analyses were performed in ADA-animals due to the impact of immunogenicity on systemic drug exposure.

### Transcriptomic and proteomic analyses

Transcriptomic and proteomic analyses were performed on matched hemi-brain tissue samples collected at study termination (n=6/sex/treatment).

## RESULTS

### ChAdu achieves dose-dependent plasma and brain exposure following treatment in aged 5XFAD mice

ChAdu was detected in both plasma and brain of aged 5XFAD mice following pilot and chronic administration, and chAdu exposure was attenuated in mice that developed ADAs (Fig. 1).

In the 4-week pilot study, a single 30 mg/kg dose of chAdu produced sustained plasma exposure through day 28 (Fig. 1B) with a model-estimated elimination half-life of 6.1 days. A one-compartment model adequately described plasma concentration-time data across pilot chAdu-treatment groups (Supplementary Fig. S1-S4; Supplementary Table S4). Weekly chAdu administration for 4 weeks resulted in peak plasma concentrations of 234.13 ± 58.25 µg/mL and 6.67 ± 0.81 µg/mL at 3 days after the fourth dose in the 30 mg/kg and 1 mg/kg groups, respectively, in ADA-mice (Fig. 1B). Terminal brain concentrations measured 7 days after the final pilot dose were 0.11 ± 0.02 pg/µg after 1 mg/kg once-weekly dosing, 5.64 ± 0.57 pg/µg after 30 mg/kg once-weekly dosing, and 1.56 ± 0.33 pg/µg after a single 30 mg/kg dose in ADA-animals (Fig. 1C).

In the chronic cohort, once-weekly chAdu administration for 17 weeks produced dose-dependent increases in plasma and brain exposure in ADA-mice. At 30 mg/kg, males exhibited higher terminal plasma concentrations than females (299.32 ± 26.86 µg/ml vs. 189.07 ± 20.39 µg/mL; *p* = 0.01), whereas females exhibited higher brain chAdu concentrations than males (19.82 ± 1.85 pg/µg vs. 12.86 ± 1.34 pg/µg; *p* < 0.0001) (Fig. 1D-E). No sex differences were observed at lower doses. Observed plasma concentrations fell within the prediction intervals generated from the pilot popPK model, supporting the adequacy of the pilot-derived model for characterizing chronic exposure across dose groups (Supplementary Fig. S1 and S5).

### Treatment-emergent ADAs are associated with reduced circulating chAdu exposure

Following 4-weeks of chAdu treatment, 4 of 21 mice (19%) screened positive for ADAs (ADA+) in terminal plasma, including two males in the 1 mg/kg weekly group and two females in the 30 mg/kg weekly group; one subject also screened ADA+ in plasma collected prior to dosing (Fig. 1F; Supplementary Table S2). ADA+ mice exhibited reduced plasma chAdu exposure compared to ADA-mice across treatment groups (Fig. 1F).

Following 17-weeks of chAdu treatment, 6 of 61 mice (10%) screened positive for ADAs, including five females and one male distributed across treatment groups (Fig. 1G, Supplementary Table S3). Across dose groups, ADA+ mice exhibited reduced plasma chAdu concentrations vs. ADA-animals when normalize to the ADA-group mean within each dose (Mann–Whitney test, *p* < 0.001; Fig. 1G).

Because ADA+ animals had markedly reduced systemic chAdu exposure, primary chronic pharmacodynamic (PD), behavioral, and multi-omic analyses were conducted in ADA-mice (Fig. 2-6). Full cohort analyses irrespective of ADA status are provided in Supplementary Figures S5-S7, and pilot PD analyses with and without ADA exclusion are summarized in Supplementary Tables S5-S7.

**Figure 2.**
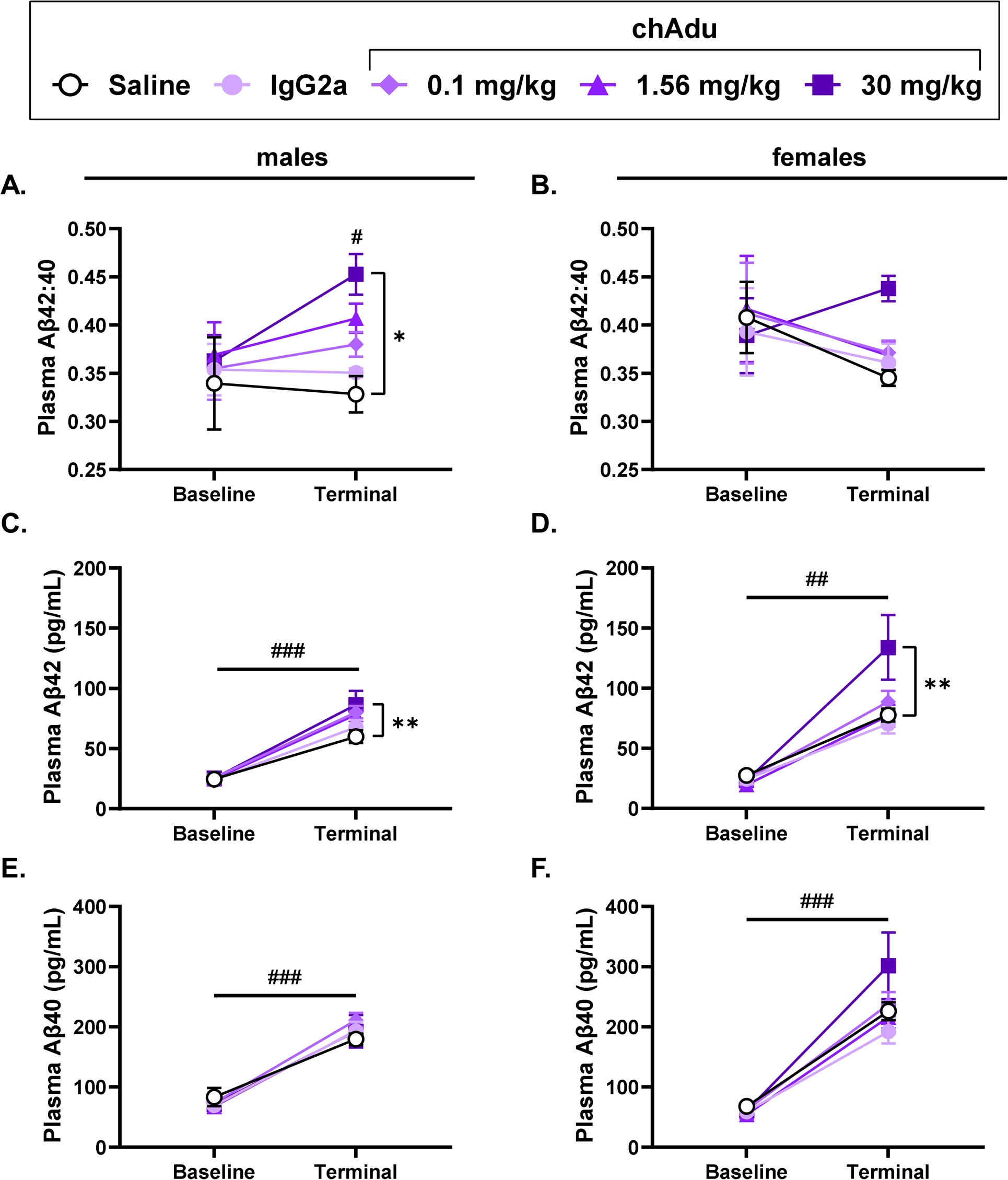
Longitudinal evaluation of plasma Aβ in aged male and female 5XFAD mice in response to chronic chAdu treatment. Data are presented only for subjects that did not develop anti-drug antibodies following once weekly 17-week treatment with murinized chimeric aducanumab (chAdu). (A–B) Plasma Aβ42:40 at baseline and terminal collection in males (A) and females (B). (C–D) Plasma Aβ42 concentrations at baseline and terminal collection in males (C) and females (D). (E–F) Plasma Aβ40 concentrations at baseline and terminal collection in males (E) and females (F). Points indicate group mean ± SEM. Data were analyzed within each sex using two-way repeated-measures ANOVA (time × treatment) followed by Tukey’s multiple comparisons test. Statistical annotations indicate: * and **, differences between treatment groups at the terminal timepoint; #, significant differences between baseline and terminal within treatment group(s). Where # is shown above a single group, the effect applies only to that group; where shown as a spanning bar, the effect applies across all groups (\**p* < 0.05, \*\**p* < 0.01; #*p* < 0.05, ##*p* < 0.01, ###*p* < 0.001).

### Plasma A**β**42:40 as a biomarker for treatment response to chAdu

Baseline plasma Aβ42:40 ratios were comparable across treatment groups within sex prior to treatment initiation (Fig. 2A-F). Plasma Aβ42:40 was evaluated longitudinally as an exploratory peripheral marker of altered Aβ dynamics following antibody treatment [23,24]. Over the 17-week treatment period, plasma Aβ42 and Aβ40 increased in aging male and female 5XFAD mice (Fig. 2C-F).

In males, plasma Aβ42:40 was unchanged from baseline to terminal in saline-and IgG-treated controls, whereas 17-week chAdu treatment produced a dose-dependent increase with elevations observed at 30 mg/kg (*p* = 0.01; Fig. 2A). In females, plasma Aβ42:40 decreased in saline-and IgG-treated controls, while the 30 mg/kg chAdu group demonstrated a non-significant upward trend relative to baseline (Fig. 2B).

Plasma Aβ42 concentrations increased from baseline to terminal across treatment groups in both sexes and were elevated in the 30 mg/kg group vs. saline-treated controls at terminal in males (86.44 ± 11.47 pg/mL vs. 59.90 ± 5.35 pg/mL, *p* = 0.0091) and females (134.0 ± 26.87 pg/mL vs. 77.82 ± 5.34 pg/mL, *p* = 0.0023) (Fig. 2C-D). No terminal differences were observed between saline-treated mice and IgG-, 0.1 mg/kg chAdu-, or 1.56 mg/kg chAdu-treated females. Plasma Aβ40 concentrations similarly increased from baseline to terminal in both sexes but did not differ between treatment groups at the terminal timepoint (Fig. 2E-F). These patterns reflect altered Aβ dynamics following anti-amyloid antibody treatment [23,25].

### Differential effects of sex on reductions of brain amyloid by both chAdu and IgG

Terminal brain Aβ40 and Aβ42 were quantified and normalized to total protein following 17 weeks of treatment in ADA-subjects and expressed as a percentage of the within-sex saline-treated control mean (Fig. 3). Absolute brain Aβ concentrations for all mice regardless of ADA status are provided in Supplementary Figure S7.

**Figure 3.**
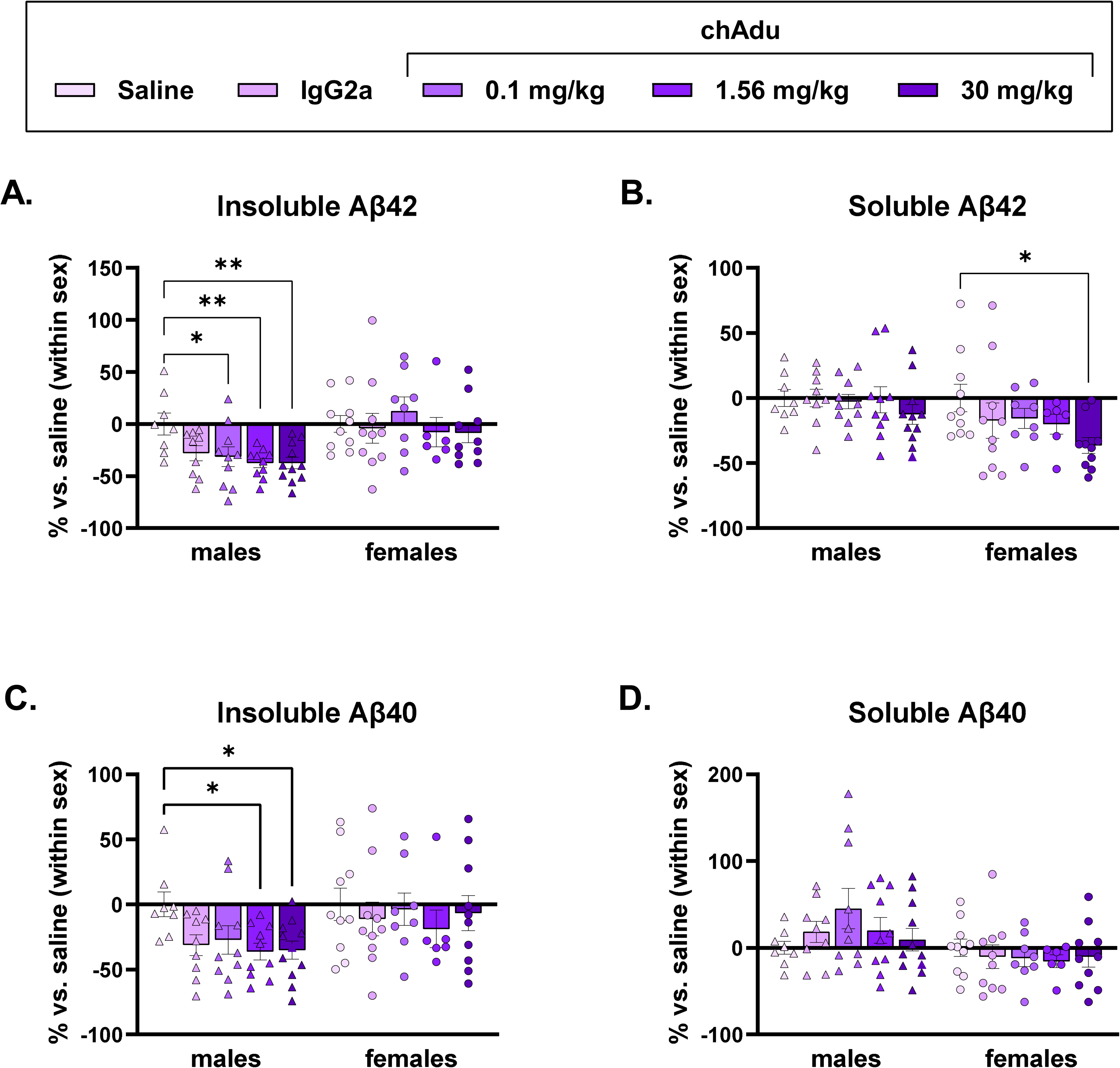
Effects of chronic chAdu treatment on brain. **A**β **in male and female 5XFAD mice.** Data are presented for only subjects that did not demonstrate anti-drug antibodies (ADA) to murinized chimeric aducanumab (chAdu) following 17-week treatment. (A) Insoluble Aβ42 concentrations in male and female mice at terminal collection. (B) Soluble Aβ42 concentrations in male and female mice at terminal collection. (C) Insoluble Aβ40 concentrations in male and female mice at terminal collection. (D) Soluble Aβ40 concentrations in male and female mice at terminal collection. Bars represent mean ± SEM, expressed as a percentage of the saline-treated group mean (set to 0%) within sex for each measure. Individual data points represent single animals (triangles, males; circles, females). Within each sex, data were analyzed by one-way ANOVA followed by Dunnett’s multiple comparisons test, with IgG-and chAdu-treated groups compared to saline-treated controls. (\**p* < 0.05, \*\**p* < 0.01).

At 30 mg/kg, chAdu treatment reduced brain Aβ42 relative to saline-treated controls. In males, insoluble Aβ42 decreased by 37.5 ± 5.65% (*p* = 0.01), while in females, soluble Aβ42 was reduced by 36.54 ± 6.13% (*p* = 0.03) (Fig. 3A-B).

In males, additional reductions in insoluble Aβ42 were observed at lower chAdu doses, including 0.1 mg/kg (31.46 ± 9.58% reduction, *p* = 0.0231) and 1.56 mg/kg (37.44 ± 4.38% reduction, *p* = 0.0055) (Fig. 3A). No significant reductions in insoluble or soluble Aβ42 were observed in females at these doses (Fig. 3A-B).

For Aβ40, reductions in the insoluble fraction were observed in males at 30 mg/kg (35.13 ± 7.0% reduction, *p* = 0.02) and 1.56 mg/kg (36.29 ± 6.43% reduction, *p* = 0.02), whereas no significant differences were observed in females (Fig. 3C). Soluble Aβ40 levels were unchanged across treatment groups in either sex (Fig. 3D).

IgG-treated males showed reductions in insoluble Aβ42 (28.12 ± 7.3% reduction, *p* = 0.06) and Aβ40 (31.29 ± 8.0% reduction, *p* = 0.06) similar in direction to chAdu-treated males, but these effects were not statistically significant (Fig. 3A,C).

### Behavioral effects of IgG and chAdu treatment

Consistent with previous reports [16], 5XFAD mice exhibited increased cumulative frailty scores vs. WT controls, with more pronounced effects in females. Frailty scores did not differ between chAdu-and IgG-treated 5XFAD mice in either sex (Fig. 4A).

**Figure 4.**
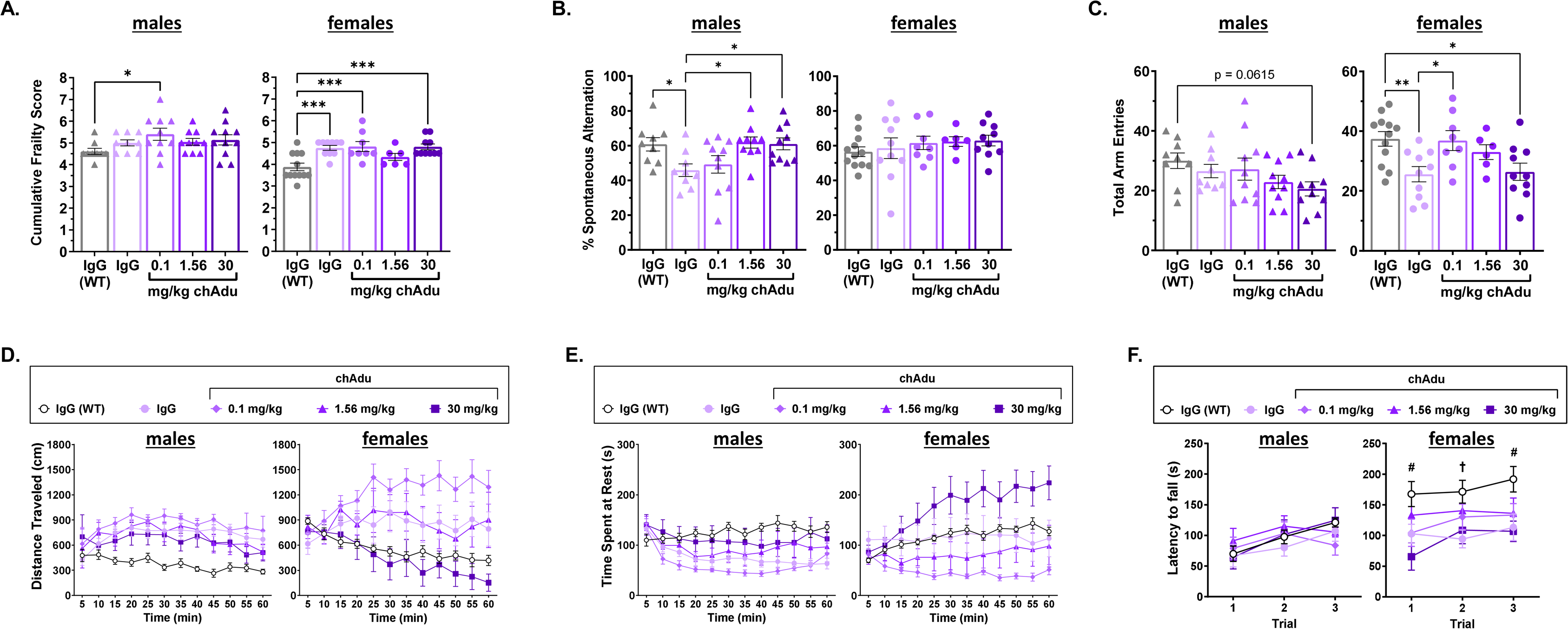
Behavioral effects of chronic chAdu treatment in aging male and female 5XFAD mice. Data are presented for only subjects that did not demonstrate anti-drug antibodies (ADA) to murinized chimeric aducanumab (chAdu) following 17-week treatment. Chronic chAdu treatment did not improve behavioral alterations in male or female 5XFAD mice with modest motor altering effects observed relative to age-and sex-matched IgG-vehicle-treated 5XFAD or IgG-treated vehicle WT controls. (A) Cumulative frailty scores in male and female mice. (B) Percent alternation in the spontaneous alternation task in male and female mice. (C) Total arm entries in the spontaneous alternation task in male and female mice. (D) Distance traveled over time in the open field test in male and female mice. (E) Time spent at rest over time in the open field test in male and female mice. (F) Rotarod performance, measured as latency to fall across three trials, in male and female mice. For panels A-C, points represent individual animals and bars indicate mean ± SEM; data were analyzed within each sex using one-way ANOVA followed by Dunnett’s multiple comparisons test, with treatment groups compared to IgG-treated WT controls. For panels D-F, points represent group mean ± SEM; data were analyzed within each sex using two-way repeated-measures ANOVA with Geisser–Greenhouse correction, followed by Dunnett’s multiple comparisons test, with treatment groups compared to IgG-treated WT controls. (\**p* < 0.05, \*\**p* < 0.01, \*\*\**p* < 0.001, \*\*\*\**p* < 0.0001, #*p* < 0.05 for 30 mg/kg chAdu vs. controls, †*p* < 0.05 for IgG-treated 5XFAD vs. controls).

In the spontaneous alternation task, IgG-treated 5XFAD males exhibited reduced percent alternation relative to WT controls (45.85 ± 3.61% vs. 60.75 ± 3.82%, *p* = 0.0392). ChAdu treatment increased alternation performance in a dose-dependent manner, with the 1.56 and 30 mg/kg groups (61.84 ± 3.16% and 60.97 ± 3.41%, respectively) reaching levels comparable to WT mice (Fig. 4B). However, these improvements coincided with reduced total arm entries (Fig. 4C). In females, percent alternation did not differ between IgG-treated 5XFAD and WT mice and was not altered by chAdu treatment. Both IgG-and high-dose chAdu-treated female 5XFAD mice exhibited reduced total arm entries vs. WT controls (Fig. 4C), indicating reduced locomotor activity.

Open field testing demonstrated locomotor alterations across treatment groups (Fig. 4D-E). Male IgG-treated 5XFAD mice exhibited increased distance traveled and reduced resting time vs. WT controls (*p* < 0.0001), consistent with the previously described hyperactivity phenotype [15]. High-dose chAdu partially attenuated this phenotype, increasing resting time (*p* = 0.0001) and modestly reducing distance traveled. Female IgG-treated 5XFAD mice also exhibited increased distance traveled vs. WT controls (*p* < 0.0001). However, unlike males, chAdu treatment produced bidirectional effects in females, with low-dose treatment increasing hyperactivity and high-dose treatment reducing locomotor activity relative to IgG-treated 5XFAD mice (Fig. 4D-E).

Rotarod performance demonstrated sex-divergent effects (Fig. 4F). All groups exhibited improved performance across repeated trials, but no treatment-associated differences were observed in males. In contrast, both IgG-and 30 mg/kg chAdu-treated female 5XFAD mice showed decreased latency to fall vs. WT controls (p < 0.01) (Fig. 4F).

### Proteomic signatures of disease progression and therapeutic response to chAdu and IgG treatment

Females exhibited more disease-associated differentially expressed proteins (DEPs) than males (2,852 vs. 2,009 DEPs; FDR < 0.05). Antibody-treated cohorts displayed more modest proteomic changes relative to the disease baseline, with female high-dose chAdu treatment producing the largest treatment-associated response (1,536 DEPs), approximately three-fold greater than IgG treatment (566 DEPs). In contrast, males exhibited relatively similar responses across chAdu and IgG treatment (Fig. 5A). Proteins upregulated in 5XFAD mice frequently shifted toward WT levels following treatment, and treatment associated protein expression changes demonstrated dose-and sex-dependent patterns (Supplementary Fig. S8).

**Figure 5:**
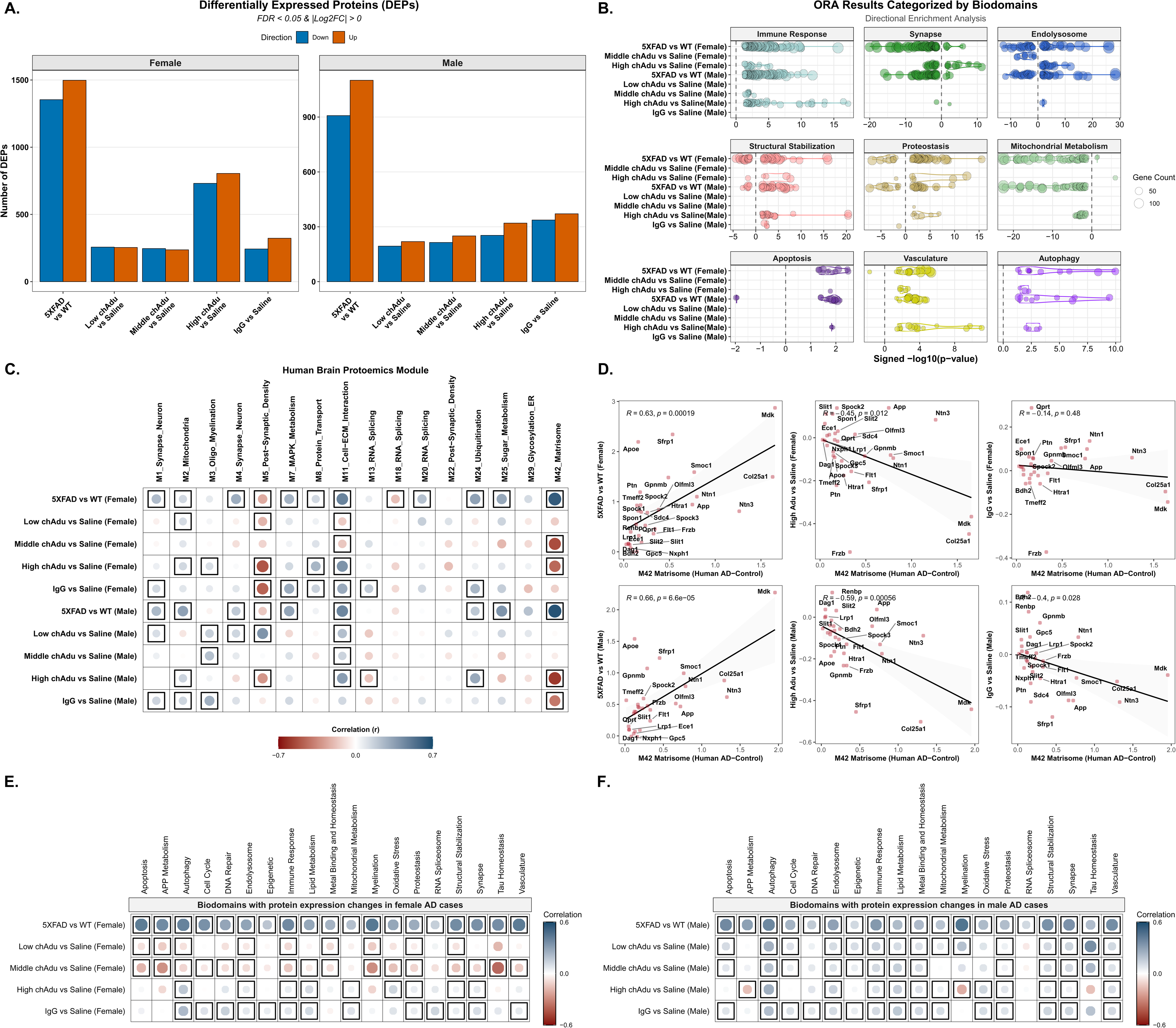
Proteomic landscape of 5XFAD pathology and translational response to chAdu immunotherapy and IgG treatment. (A) Differential expression profiles: Bar plots illustrating the total count of significantly up-or down-regulated proteins (FDR <0.05) across experimental groups for females (left) and males (right). Comparisons include the baseline 5XFAD genotype effect (IgG-5XFAD vs. IgG-WT) and the therapeutic effects of low-, middle-, and high-dose chAdu or IgG (relative to saline-5XFAD controls). (B) Directional Over-Representation Analysis (ORA) by AD Biological domains. Dot plot summarizing Over-Representation Analysis (ORA) of differentially expressed proteins categorized into 19 Alzheimer’s disease-related biological domains based on their associated GO identifiers, highlighting pathway level alterations relevant to Alzheimer’s disease. The x-axis represents the signed-log10(p-value), where positive values indicate enrichment in upregulated genes and negative values indicate enrichment in downregulated genes. Circle size represents the gene count (number of genes) associated with each enriched term. (C) Correlation with human AD proteomic modules10. Heatmap showing Pearson correlation coefficients between protein expression change in mouse groups (log fold change of IgG treated 5XFAD mice relative to sex-matched IgG treated WT littermates, and of treated 5XFAD mice relative to sex-matched saline treated 5XFAD controls) and protein expression changes in human AD subjects (log fold change for cases versus controls). Blue and red circles represent positive and negative correlations, respectively; circle size and color intensity reflect the correlation magnitude. Significant associations (*p* < 0.05) are highlighted with black frames. (D) Matrisome-specific translational alignment. Scatter plots correlating the expression changes of proteins within the M42_Matrisome module. Scatter plots showing correlations between expression changes of M42_Matrisome module proteins in human AD versus 5XFAD mice (left column) and human AD versus high dose chAdu or IgG treated 5XFAD mice (right column). These plots illustrate the degree to which treatment reverses or maintain the disease-associated matrisome signature. (E-F) Translational conservation across biological domains. Heatmap showing the correlation between mouse proteomic signatures and human AD protein signatures across 19 biological domains for (E) female and (F) and males. Circles represent positive (blue) or negative (red) correlations; circle size and color intensity reflect correlation strength, with significant correlations (p < 0.05) framed. Note: Throughout all panels, “Low chAdu”, “Middle chAdu”, and “High chAdu” refer to 5XFAD mice treated with low-, middle-, or high-dose aducanumab, respectively; “IgG” and “Saline” denote 5XFAD mice treated with the IgG isotype control or saline vehicle.

**Figure 6:**
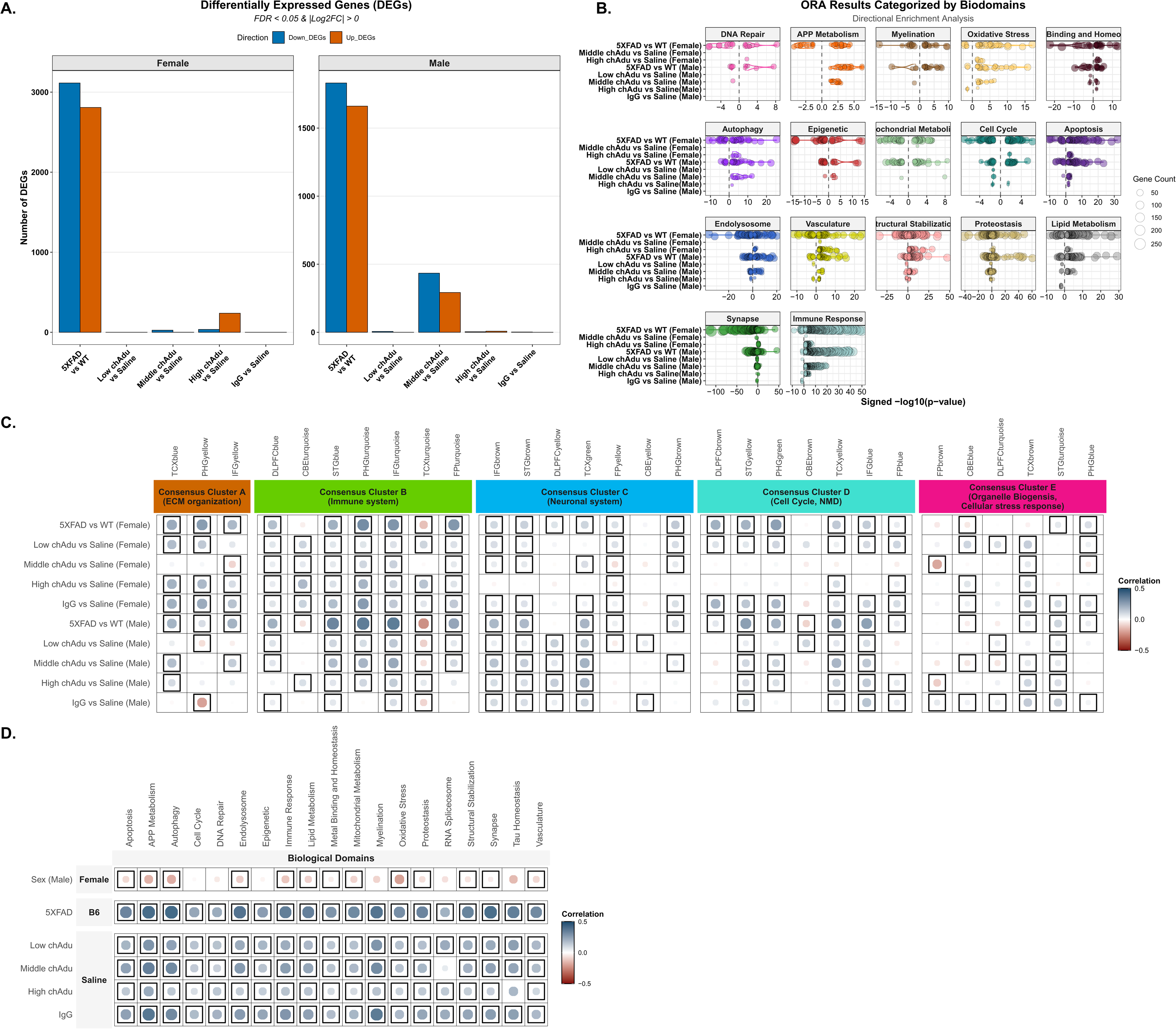
Transcriptomic landscape of 5XFAD pathology and translational response to chAdu treatment. (A) Differential gene expression across experimental cohorts. Bar plots displaying the total number of significantly up-or down-regulated genes (FDR <0.05) in IgG treated 5XFAD mice (relative to IgG treated WT) and in low-, middle-, and high-dose chAdu or IgG treated 5XFAD mice (relative to saline treated 5XFAD) for both females (left) and males (right). (B) Directional Over-Representation Analysis (ORA) by AD Biodomains. Dot plot summarizing the functional enrichment of differentially expressed genes categorized into 19 Alzheimer’s Disease-related biological domains. The plot highlights pathway-level alterations, showing the shift in transcriptomic signatures associated with neuroinflammation, synaptic function, and metabolic pathways. The x-axis represents the signed-log10(p-value, where positive values indicate enrichment in upregulated genes and negative values indicate enrichment in downregulated genes. Circle size represents the gene count (number of genes) associated with each enriched term. (C) Correlation with human AMP-AD co-expression modules11. Heatmap correlating mouse transcriptomic signatures with 30 human AD co-expression modules. Human modules are grouped into five previously identified consensus clusters describing the major functional groups of AD-related alterations11. The heatmap shows Pearson correlation coefficients between gene expression change in mouse groups (log fold change of IgG treated 5XFAD mice relative to sex-matched IgG treated WT, and of chAdu-treated 5XFAD mice relative to sex-matched saline treated 5XFAD controls) and gene expression differences in human AD subjects (log fold change for cases versus controls). Circles denote positive (blue) or negative (red) correlations; color intensity and circle size reflect the correlation magnitude, with significant correlations (p < 0.05) indicated by frames. (D) Correlation between transcriptomic effects of sex (male), 5XFAD genotype, and low, middle, or high dose chAdu and IgG treatment and human AD transcriptomic signatures across 19 AD associated biological domains. Circles represent positive (blue) or negative (red) correlations; circle size and color intensity reflect correlation strength, with significant correlations (p < 0.05) framed. Note: Throughout all panels, “Low chAdu”, “Middle chAdu”, and “High chAdu” refer to 5XFAD mice treated with low-, middle-, or high-dose aducanumab, respectively; “IgG” and “Saline” denote 5XFAD mice treated with the IgG isotype control or saline vehicle.

### Functional enrichment and translational analyses of proteomic changes

GO enrichment analysis identified Immune Response, Apoptosis, Autophagy, Structural Stabilization, and Vasculature biodomains among proteins upregulated in 5XFAD mice relative to WT in both sexes, whereas proteins downregulated in 5XFAD mice were enriched for Synapse and Mitochondrial Metabolism pathways. Proteostasis and Endolysosome pathways were enriched across both upregulated and downregulated protein sets. High-dose chAdu modulated synapse-related pathways in females and upregulated immune response-related proteins in both sexes (Fig. 5B).

Male and female 5XFAD mice exhibited significant positive correlations with several human AD-associated proteomic modules, including Synapse/Neuron, Mitochondria, MAPK Metabolism, Cell–ECM Interaction, and Matrisome (Fig. 5C). ChAdu and IgG partially reversed these signatures. Female cohorts demonstrated significant negative correlations with the Cell–ECM Interaction module at low and middle doses and with the Matrisome module at middle and high doses. Similar matrisome-related reversals were observed in males receiving high-dose chAdu or IgG (Fig. 5C). Still, positive correlations persisted across several modules at all chAdu doses. Analysis of the M42_Matrisome module identified Smoc1, Apoe, App, and Mdk among the proteins contributing to treatment-associated shifts (Fig. 5D).

Male and female saline-treated 5XFAD mice exhibited significant positive correlations across all AD-associated biological domains (Fig. 5E-F). In females, low-and middle-dose chAdu treatment elicited significant negative correlations with most biological domains (Fig. 5E), whereas male treatment groups generally retained positive correlations regardless of dose, with only high-dose chAdu eliciting significant negative correlations in two domains (Fig. 5F).

### Differential gene expression and functional enrichment across AD domains

In female 5XFAD mice, substantially more genes were differentially expressed relative to WT controls than in males (5,922 vs. 3,492 differentially expressed genes (DEGs); FDR < 0.05), paralleling the proteomic analyses and indicating a stronger baseline molecular response to amyloid pathology. In contrast, chAdu and IgG treatment produced relatively few transcriptomic alterations, with significant DEGs detected only in middle-dose chAdu-treated males and high-dose chAdu-treated females (Fig. 6A).

In both sexes, genes associated with 5XFAD genotype were enriched for GO terms related to Immune Response, Synapse, Oxidative Stress, Autophagy, Vasculature, and Mitochondrial Metabolism. Immune Response terms were predominantly enriched among upregulated genes, whereas Synapse-related terms were enriched among downregulated genes in both sexes, indictive of the classic AD molecular signature of neuroinflammation coupled with synaptic loss (Fig. 6B).

Treatment-associated DEGs demonstrated enrichment of Immune Response, Apoptosis, Structural Stabilization, Vasculature, and Oxidative Stress domains among upregulated genes in chAdu-treated 5XFAD mice of both sexes. Male-specific responses additionally included enrichment of APP metabolism-associated pathways among upregulated genes. The Synapse endophenotype exhibited modest but significant enrichment across both upregulated and downregulated gene sets in both sexes (Fig. 6B).

### Correlation of mouse transcriptomic signatures with human AD consensus clusters

In both sexes, 5XFAD mice exhibited significant positive correlations across all five human AMP-AD consensus clusters, indicating broad recapitulation of human AD-associated transcriptomic signatures (Fig. 6C). Although chAdu and IgG treatment generally maintained positive correlations with human AD modules, several dose-and sex-specific reversals were identified. In females, middle-dose chAdu treatment demonstrated significant negative correlations with the ECM-associated IFGyellow module, whereas middle-and high-dose treatment negatively correlated with the synapse-associated FPyellow module. In males, low-dose chAdu and IgG treatment negatively correlated with the ECM-associated PHGyellow module, and low-dose chAdu demonstrated reversal of the synapse-associated FPyellow module (Fig. 6C).

All experimental cohorts, including baseline 5XFAD and treatment groups, exhibited significant positive correlations across AD-associated biological domains (Fig. 6D), indicating antibody treatment did not globally reverse disease-associated transcriptomic endophenotypes.

## DISCUSSION

The present study provides an integrated evaluation of chronic chAdu treatment, IgG-associated effects, sex-dependent responses, and post-treatment molecular signatures in aged 5XFAD mice. ChAdu altered plasma and brain Aβ dynamics, although brain Aβ lowering was modest relative to the extensive amyloid burden characteristic of aged 5XFAD mice, which may be challenging to overcome at any dose. Dose selection was informed by prior Tg2576 efficacy studies [6], allowing evaluation of whether previously reported pharmacologic effects translated to aged 5XFAD mice with established pathology. Importantly, biologically meaningful effects were also observed with the IgG2aκ isotype control antibody, emphasizing the need to distinguish IgG-associated mechanisms from antigen-specific therapeutic effects in preclinical immunotherapy studies.

ChAdu treatment increased plasma Aβ42 and plasma Aβ42:40 ratio while producing modest reductions in brain Aβ species that were directionally consistent with antibody-mediated plaque engagement. However, these effects were less robust than previously reported in Tg2576 mice, suggesting pharmacodynamic expectations derived from one amyloidogenic model may not translate quantitatively to aged 5XFAD mice. In males, both chAdu and IgG treatment produced similar trends toward reduced insoluble Aβ species, supporting a potential contribution of Fc receptor-mediated or immune-modulatory mechanisms independent of high-affinity Aβ binding [25–29].

Although aducanumab, lecanemab, and donanemab target distinct Aβ species, they share a common IgG backbone [30], which itself can influence AD-relevant biology through Fc receptor signaling, microglial activation, and modulation of peripheral and central Aβ dynamics independent of high-affinity antigen binding [25]. Intravenous immunoglobulin (IVIG) preparations similarly exhibit anti-inflammatory and AD-relevant biological effects [26,27], suggesting IgG-associated mechanisms may contribute to therapeutic and adverse effects observed with amyloid-targeting antibodies [25] which are consistent with our results.

A prominent feature of the current study was the presence of sex-dependent treatment responses across biochemical, molecular, and behavioral measures. Males exhibited greater reductions in insoluble Aβ species and stronger shifts in Aβ-associated proteomic modules, whereas females required higher brain chAdu exposure to demonstrate reductions in soluble Aβ42 or convergent proteomic responses. These findings align with known sex differences in 5XFAD pathology and broader evidence that sex influences amyloid pathology, immune response, and therapeutic outcomes following Aβ-targeting immunotherapy [17]. The observed sex-dependent effects likely reflect differences in brain disposition, Fc-mediated biology, immunogenicity, or downstream pharmacodynamic mechanisms rather than systemic chAdu PK. These findings underscore the need for rigorous consideration of sex as a biological variable in preclinical and clinical evaluations of immunotherapies.

Treatment-emergent ADA incidence ranged from 10-19% across the pilot and chronic studies, and ADAs were associated with reduced plasma chAdu exposure, consistent with enhanced drug clearance in mice mounting an anti-drug immune response. Similar exposure reductions have been reported clinically for therapeutic antibodies [31]. Although overall ADA incidence was low, the predominance of ADA+ females further highlights the interaction between sex, immune reactivity, and therapeutic exposure.

Multi-omics analyses demonstrated that both chAdu and IgG treatment modulated Aβ-associated proteomic networks. In particular, Module 42, which contains Aβ-and plaque-associated proteins, exhibited significant negative correlations in high-dose chAdu-and IgG-treated males relative to saline controls. Additional synaptic and signaling pathways were altered following antibody treatment, particularly in females, where postsynaptic density-associated Module 5 demonstrated significant negative correlations across all antibody-treated groups, including IgG controls. These findings further support biologically meaningful IgG-associated effects independent of antigen-specific Aβ targeting.

Despite measurable amyloid lowering, several disease-associated pathways remained unaltered following treatment. Mitochondrial Complex I subunits such as Ndufa7, as well as synaptic receptor and signaling targets including Gabra1, Chrna4, and Scn2a, remained dysregulated following antibody exposure. Many of these pathways are currently under investigation as therapeutic targets [32], emphasizing amyloid lowering alone may not sufficiently address broader drivers of AD pathobiology.

Several model-related considerations are important when interpreting these findings. The rapid and early plaque accumulation characteristic of 5XFAD mice, particularly in females, likely limits the detectable biochemical impact of treatment once pathology is well established. Although dose-dependent proteomic changes were identified, brain Aβ alterations measured biochemically were less robust at lower doses, suggesting plaque deposition may outpace antibody-mediated clearance in this model. These findings support continued development of next-generation models such as hAbetaSAA mice (JAX #034711), which exhibit more gradual and physiologically relevant plaque accumulation without the hyperactivity confounds common in transgenic models [15,33]. Such models may better capture the therapeutic window within which immunotherapies can meaningfully influence Aβ accumulation and downstream processes.

Taken together, chronic chAdu treatment produced measurable but modest effects on plasma and brain Aβ in aged 5XFAD mice while revealing pronounced sex-dependent and IgG-associated molecular responses. These findings highlight the importance of considering Fc-mediated immune modulation and sex when interpreting antibody-based interventions. More broadly, this work demonstrates the value of integrated preclinical pipelines combining biochemical, behavioral, PK, ADA, transcriptomic, and proteomic analyses for identifying disease mechanisms that remain insufficiently addressed by amyloid-targeting immunotherapy alone.

## Acknowledgements

This work was supported by the National Institutes of Health, National Institute on Aging (U54 AG054345). The authors are grateful for the support of the University of Pittsburgh Preclinical Phenotyping Core and the Genome Technologies Core at The Jackson Laboratory. Mass spectrometry analyses supporting pharmacokinetic assessment of chAdu were performed by the Indiana University School of Medicine Center for Proteome Analysis. Acquisition of instrumentation used for these studies was supported by the Indiana University Precision Health Initiative, and this work was also supported, in part, by the Indiana Clinical and Translational Sciences Institute (UL1TR002529) and the IU Simon Comprehensive Cancer Center Support Grant (P30CA082709).

## Conflicts of Interest

Jeffrey L. Dage (JLD) is an inventor on patents or patent applications assigned to Eli Lilly and Company relating to the assays, methods, reagents and / or compositions of matter for P-tau assays and Aβ targeting therapeutics. JLD has/is served/serving as a consultant or on advisory boards for Eisai, Abbvie, Genotix Biotechnologies Inc, Gates Ventures, Syndeio Biosciences, Dolby Family Ventures, Karuna Therapeutics, Alzheimer’s Disease Drug Discovery Foundation, AlzPath Inc., Cognito Therapeutics, Inc., Eli Lilly and Company, Prevail Therapeutics, Neurogen Biomarking, Spear Bio, Rush University, Washington University in St. Louis, University of Kentucky, Tymora Analytical Operations, MindImmune Therapeutics, Inc, Early is Good, Roche and Quanterix. JLD has received research support from ADx Neurosciences, Fujirebio, Roche Diagnostics and Eli Lilly and Company in the past two years. JLD has received speaker fees from Eli Lilly and Company and LabCorp. JLD is a founder and advisor for Monument Biosciences and Dage Scientific LLC. JLD has stock or stock options in Eli Lilly and Company, Genotix Biotechnologies, MindImmune Therapeutics Inc., AlzPath Inc., Neurogen Biomarking, and Monument Biosciences.

All other authors declare that they have no known competing financial interests or personal relationships that could have appeared to influence the work reported in this paper.

## Funding

The authors are supported by funding from the National Institutes of Health, National Institute on Aging U54 AG05434503.

## Consent Statement

No human subjects were used in these studies, and, therefore, consent was not necessary.

ZAC, S-PW, SKQ, MS, BTL, ALO, GWC, PRT, and SJSR contributed to study conceptualization and design. KAH, EHD, SKQ, JLD, DMD, NTS, PRT, and SJSR contributed to methodology development and experimental workflows. KAH, EHD ZAC, GJL, S-PW, UN, and DMD performed experiments and collected data. KAH, RSP, SKQ, AN, NBC, LdS, GWC, and SJSR contributed to data analysis and interpretation. KAH, RSP, EHD, ZAC, GJL, S-PW, and DMD contributed to data organization. KAH, RSP, SKQ, AN, and SJSR generated and edited figures and tables. KAH, RSP, EHD, AN, GWC, and SJSR contributed to writing the original draft of the manuscript. KAH, SKQ, AN, LdS, JLD, NTS, MS, BTL, ALO, GWC, PRT, and SJSR contributed to review and editing of the manuscript. KAH, EHD, GJL, and S-PW contributed to project administration. SKQ, LdS, JLD, NTS, MS, BTL, ALO, GWC, PRT, and SJSR provided supervision. MS, BTL, ALO, GWC, PRT, and SJSR acquired funding for the study.

## Abbreviations

chAdu: murine chimeric aducanumab
ADA: anti-drug antibody
popPK: population PK
MODEL-AD: Model Organism Development and Evaluation for Late-onset Alzheimer’s Disease
PTC: Preclinical Testing Core.

## Notes

https://doi.org/10.7303/syn74927054

https://panoramaweb.org/Indiana%20U%20-%20Proteome%20Analysis/Targeted%20quantitation%20of%20chimeric%20aducanumab%20in%20mouse%20plasma%20and%20cortex/project-begin.view

https://panoramaweb.org/Indiana%20U%20-%20Proteome%20Analysis/Reproducability%20and%20repeatability%20Aducanumab%20Assay/project-begin.view

## REFERENCES

[1] Sukoff Rizzo SJ, Masters A, Onos KD, Quinney S, Sasner M, Oblak A, et al. Improving preclinical to clinical translation in Alzheimer’s disease research. Alzheimer’s and Dementia: Translational Research and Clinical Interventions 2020;6. 10.1002/trc2.12038.

[2] Kim CK, Lee YR, Ong L, Gold M, Kalali A, Sarkar J. Alzheimer’s Disease: Key Insights from Two Decades of Clinical Trial Failures. Journal of Alzheimer’s Disease 2022;87:83–100. 10.3233/JAD-215699.

[3] Torres A, Camargo L, López N. Aducanumab: A look two years after its approval. Biomedica 2024;44:42–6. 10.7705/BIOMEDICA.6967.

[4] Heidebrink JL, Paulson HL. Lessons Learned from Approval of Aducanumab for Alzheimer’s Disease. Annu Rev Med 2024;75:99–111. 10.1146/annurev-med-051022-043645.

[5] Budd Haeberlein S, Aisen PS, Barkhof F, Chalkias S, Chen T, Cohen S, et al. Two Randomized Phase 3 Studies of Aducanumab in Early Alzheimer’s Disease. J Prev Alzheimers Dis 2022;9:197–210. 10.14283/jpad.2022.30.

[6] Sevigny J, Chiao P, Bussière T, Weinreb PH, Williams L, Maier M, et al. The antibody aducanumab reduces Aβ plaques in Alzheimer’s disease. Nature 2016;537:50–6. 10.1038/nature19323.

[7] Thussu S, Naidu A, Manivannan S, Grossberg GT. Profiling aducanumab as a treatment option for Alzheimer’s disease: an overview of efficacy, safety and tolerability. Expert Rev Neurother 2024;24:1045–53. 10.1080/14737175.2024.2402058.

[8] Salloway S, Pain A, Lee E, Papka M, Ferguson MB, Wang H, et al. TRAILBLAZER-ALZ 4: A phase 3 trial comparing donanemab with aducanumab on amyloid plaque clearance in early, symptomatic Alzheimer’s disease. Alzheimer’s and Dementia 2025;21. 10.1002/alz.70293.

[9] Qi X, Nizamutdinov D, Yi SS, Wu E, Huang JH. Disease Modifying Monoclonal Antibodies and Symptomatic Pharmacological Treatment for Alzheimer’s Disease. Biomedicines 2024;12. 10.3390/biomedicines12112636.

[10] Heneka MT, Morgan D, Jessen F. Passive anti-amyloid β immunotherapy in Alzheimer’s disease—opportunities and challenges. The Lancet 2024;404:2198–208. 10.1016/S0140-6736(24)01883-X.

[11] Tonegawa-Kuji R, Hou Y, Hu B, Lorincz-Comi N, Pieper AA, Tousi B, et al. Efficacy and safety of passive immunotherapies targeting amyloid beta in Alzheimer’s disease: A systematic review and meta-analysis. PLoS Med 2025;22. 10.1371/journal.pmed.1004568.

[12] Digma LA, Winer JR, Greicius MD. Substantial Doubt Remains about the Efficacy of Anti-Amyloid Antibodies. Journal of Alzheimer’s Disease 2024;97:567–72. 10.3233/JAD-231198.

[13] Andrews D, Ducharme S, Chertkow H, Sormani MP, Collins DL. The higher benefit of lecanemab in males compared to females in CLARITY AD is probably due to a real sex effect. Alzheimer’s and Dementia 2025;21. 10.1002/alz.14467.

[14] Oblak AL, Forner S, Territo PR, Sasner M, Carter GW, Howell GR, et al. Model organism development and evaluation for late-onset Alzheimer’s disease: MODEL-AD. Alzheimer’s and Dementia: Translational Research and Clinical Interventions 2020;6. 10.1002/TRC2.12110.

[15] Oblak AL, Lin PB, Kotredes KP, Pandey RS, Garceau D, Williams HM, et al. Comprehensive Evaluation of the 5XFAD Mouse Model for Preclinical Testing Applications: A MODEL-AD Study. Front Aging Neurosci 2021;13. 10.3389/fnagi.2021.713726.

[16] Oblak AL, Cope ZA, Quinney SK, Pandey RS, Biesdorf C, Masters AR, et al. Prophylactic evaluation of verubecestat on disease-and symptom-modifying effects in 5XFAD mice. Alzheimer’s and Dementia: Translational Research and Clinical Interventions 2022;8. 10.1002/trc2.12317.

[17] Percie du Sert N, Hurst V, Ahluwalia A, Alam S, Avey MT, Baker M, et al. The ARRIVE guidelines 2.0: Updated guidelines for reporting animal research. PLoS Biol 2020;18:e3000410. 10.1371/journal.pbio.3000410.

[18] Doud EH, Hansen K, Haynes KA, Eldridge K, Arrivalagan J, Charbe N, et al. Analytical development and application of a targeted liquid chromatography-tandem mass spectrometry assay for chimeric aducanumab. MAbs 2025;17. 10.1080/19420862.2025.2537118.

[19] Carr SA, Abbatiello SE, Ackermann BL, Borchers C, Domon B, Deutsch EW, et al. Targeted peptide measurements in biology and medicine: Best practices for mass spectrometry-based assay development using a fit-for-purpose approach. Molecular and Cellular Proteomics 2014;13:907–17. 10.1074/mcp.M113.036095.

[20] Chen YQ, Pottanat TG, Carter QL, Troutt JS, Konrad RJ, Sloan JH. Affinity capture elution bridging assay: A novel immunoassay format for detection of anti-therapeutic protein antibodies. J Immunol Methods 2016;431:45–51. 10.1016/j.jim.2016.02.008.

[21] Ferrero J, Williams L, Stella H, Leitermann K, Mikulskis A, O’Gorman J, et al. First-in-human, double-blind, placebo-controlled, single-dose escalation study of aducanumab (BIIB037) in mild-to-moderate Alzheimer’s disease. Alzheimer’s and Dementia: Translational Research and Clinical Interventions 2016;2:169–76. 10.1016/j.trci.2016.06.002.

[22] Casali B, Landreth G. Aβ Extraction from Murine Brain Homogenates. Bio Protoc 2016;6. 10.21769/bioprotoc.1787.

[23] Hayato S, Takenaka O, Sreerama Reddy SH, Landry I, Reyderman L, Koyama A, et al. Population pharmacokinetic pharmacodynamic analyses of amyloid positron emission tomography and plasma biomarkers for lecanemab in subjects with early Alzheimer’s disease. CPT Pharmacometrics Syst Pharmacol 2022;11:1578–91. 10.1002/psp4.12862.

[24] Pontecorvo MJ, Lu M, Burnham SC, Schade AE, Dage JL, Shcherbinin S, et al. Association of Donanemab Treatment With Exploratory Plasma Biomarkers in Early Symptomatic Alzheimer Disease. JAMA Neurol 2022;79:1250. 10.1001/jamaneurol.2022.3392.

[25] Loeffler DA. Antibody-Mediated Clearance of Brain Amyloid-β: Mechanisms of Action, Effects of Natural and Monoclonal Anti-Aβ Antibodies, and Downstream Effects. J Alzheimers Dis Rep 2023;7:873–99. 10.3233/ADR-230025.

[26] Fei Z, Pei R, Pan B, Ye S, Zhang R, Ma L, et al. Antibody Assay and Anti-Inflammatory Function Evaluation of Therapeutic Potential of Different Intravenous Immunoglobulins for Alzheimer’s Disease. Int J Mol Sci 2023;24. 10.3390/ijms24065549.

[27] Kile S, Au W, Parise C, Sohi J, Yarbrough T, Czeszynski A, et al. Reduction of Amyloid in the Brain and Retina After Treatment With IVIG for Mild Cognitive Impairment. Am J Alzheimers Dis Other Demen 2020;35. 10.1177/1533317519899800.

[28] Abraham CR, Schneeberger A, III AFS, Hoffmann T, Fischer A, Trembleau S. AD04 – modifying Alzheimer’s Disease by modulation of neuroinflammation. Alzheimer’s & Dementia 2024;20. 10.1002/alz.095657.

[29] Haaland B, Dickson SP, Santana AF, Tanzi RE, Dubois B, Peters O, et al. Efficacy of AD04, an aluminum-based vaccine adjuvant, in patients with early Alzheimer’s disease: Post hoc analysis of AFF006 (NCT01117818), a proof-of-concept, phase 2 randomized controlled trial. Journal of Alzheimer’s Disease 2025;108:234–42. 10.1177/13872877251375985.

[30] Zhang Y, Chen H, Li R, Sterling K, Song W. Amyloid β-based therapy for Alzheimer’s disease: challenges, successes and future. Signal Transduct Target Ther 2023;8. 10.1038/s41392-023-01484-7.

[31] Mullins GR, Ardayfio P, Gueorguieva I, Anglin G, Bailey J, Chua L, et al. Donanemab immunogenicity in participants with early symptomatic Alzheimer’s disease. Alzheimer’s & Dementia: Translational Research & Clinical Interventions 2025;11. 10.1002/trc2.70149.

[32] Cary GA, Wiley JC, Carter GW, Levey AI. Beyond the streetlight: a TREAT AD perspective on where to find new Alzheimer’s targets. Alzheimer’s & Dementia 2026;22. 10.1002/alz.71142.

[33] Xia D, Lianoglou S, Sandmann T, Calvert M, Suh JH, Thomsen E, et al. Novel App knock-in mouse model shows key features of amyloid pathology and reveals profound metabolic dysregulation of microglia. Mol Neurodegener 2022;17. 10.1186/s13024-022-00547-7.

