## Supplementary Tables for "Molecular mechanisms underlying amyloid lowering by aducanumab: differential and comparative effects of sex and IgG reveal the post-treatment disease brain"

**LIST OF SUPPLEMENTARY TABLES**


| **Table** | **Description** |
|---|---|
| Table S1 | Chronic study mice excluded from terminal analysis based on a priori exclusion criteria or attrition. |
| Table S2 | Pilot study group sizes and anti-drug antibody (ADA) incidence. |
| Table S3 | Chronic study group sizes across analysis stages. |
| Table S4 | Final population pharmacokinetic model parameter estimates for chimeric aducanumab in aged 5XFAD mice. |
| Table S5 | Pilot study plasma Aβ concentrations in all mice regardless of ADA status. |
| Table S6 | Pilot study plasma Aβ concentrations in ADA-negative mice only. |
| Table S7 | Pilot study brain Aβ concentrations in all study mice and the ADA-negative subset. |

**Table S1. Chronic study mice excluded from terminal analysis based on *a priori* exclusion criteria or attrition.**

| Subject ID | Microchip ID | Genotype | Sex | Treatment Group | Reason for Exclusion |
|---|---|---|---|---|---|
| 1 | 290034992 | 5XFAD | F | Saline | >20% weight loss and seizures; deceased before Week 12 |
| 22 | 289916913 | 5XFAD | F | 1.56 mg/kg chAdu | Found dead before Week 12 |
| 50 | 289916010 | WT | M | IgG | Drug quantified in terminal plasma |
| 51 | 289916512 | WT | M | IgG | Drug detected in terminal plasma |
| 56 | 290035400 | 5XFAD | M | IgG | Drug detected in terminal plasma |
| 65 | 289916091 | WT | M | IgG | Drug quantified in terminal brain and plasma |
| 82 | 289916250 | 5XFAD | M | IgG | Drug quantified in terminal plasma |
| 119 | 290035319 | 5XFAD | F | 1.56 mg/kg chAdu | >20% weight loss; deceased before Week 8 |
| 140 | 54828-2-6 | 5XFAD | M | Saline | Drug quantified in terminal brain and plasma |
| 144 | 291019058 | 5XFAD | M | Saline | Drug detected in terminal plasma |

Mice were excluded prior to unblinding if they met predefined criteria, including (i) mortality or humane euthanasia due to health decline (e.g., >20% body weight loss or seizures) before study completion, or (ii) evidence of misdosing, defined as detectable or quantifiable drug levels in terminal plasma or brain in animals assigned to saline- or IgG-treated control groups. All exclusions were applied consistently across groups and were independent of study outcomes.

**Table S2. Pilot study group sizes and anti-drug antibody (ADA) incidence.**

| **Treatment Groups** | **Mice per group at study start** M \| F | **ADA+ at terminal** M \| F | **ADA+ pre-treatment** M \| F | **Mice per group after ADA+ exclusions** M \| F |
|---|---|---|---|---|
| 30 mg/kg IgG weekly | 3 \| 4 | 0 \| 0 | 0 \| 0 | 3 \| 4 |
| 1 mg/kg chAdu weekly | 3 \| 4 | 2 \| 0 | 0 \| 0 | 1 \| 4 |
| 30 mg/kg chAdu weekly | 3 \| 4 | 0 \| 2 | 0 \| 1 | 3 \| 2 |
| 30 mg/kg chAdu single | 3 \| 4 | 0 \| 0 | 0 \| 0 | 3 \| 4 |

Group sizes are shown by sex (M, male; F, female). No study attrition or a priori exclusions occurred in the pilot study. ADA status was assessed at terminal collection and in pre-treatment (baseline) plasma samples. One female mouse in the 30 mg/kg chAdu weekly group was ADA+ at both baseline and terminal.

**Table S3. Chronic study group sizes across analysis stages.**

| **Treatment Groups** | **Mice per group at study start** M \| F | **Mice per group after attrition** M \| F | **Mice per group after *a priori* exclusions** M \| F | **Mice per group after ADA+ exclusions** M \| F |
|---|---|---|---|---|
| Saline | 10 \| 11 | 10 \| 10 | 8 \| 10 | 8 \| 10 |
| IgG | 11 \| 10 | 11 \| 10 | 9 \| 10 | 9 \| 10 |
| 0.1 mg/kg chAdu | 10 \| 10 | 10 \| 10 | 10 \| 10 | 10 \| 8 |
| 1.56 mg/kg chAdu | 11 \| 10 | 11 \| 8 | 11 \| 8 | 10 \| 6 |
| 30 mg/kg chAdu | 11 \| 11 | 11 \| 11 | 11 \| 11 | 11 \| 10 |
| IgG (WT) | 12 \| 12 | 12 \| 12 | 9 \| 12 | 9 \| 12 |

Group sizes (male | female) are shown at study initiation, after attrition (mortality or humane endpoint prior to study completion), after application of a priori exclusion criteria (including removal of misdosed animals with detectable or quantifiable drug levels in control groups), and after exclusion of ADA+ mice for primary analyses.

**Table S4. PopPK model parameter estimates for chAdu in aged 5XFAD mice.**

| **Parameter** | **Estimate (%RSE)** |
|---|---|
| Ka ($day^{-1}$) | 1.0 |
| CL/F* (L/day/kg) | 0.03 (4.3) |
| V/F* (L/kg) | 0.23 (7.0) |
| BSV on CL/F* (%CV) | 80.2 |
| BSV on V/F* (%CV) | 46.3 |
| Proportional residual error (%) | 23.6 |

Final population pharmacokinetic model parameter estimates for chimeric aducanumab in aged 5XFAD mice. Estimates were obtained by FOCEI in nlmixr2 from pooled chAdu pilot data (N = 21 mice; 30 mg/kg single dose, 1 mg/kg once weekly, 30 mg/kg once weekly). Ka was fixed at 1 $day^{-1}$. CL/F and V/F are apparent parameters expressed per kilogram of body weight. BSV, between-subject variability expressed as approximate coefficient of variation; %RSE, relative standard error of the typical value.

**Table S5. Pilot study plasma Aβ concentrations.**

| | | Weekly 30 mg/kg IgG (3M, 4F) | | Weekly 1 mg/kg chAdu (3M, 4F) | | Weekly 30 mg/kg chAdu (3M, 4F) | | Single 30 mg/kg chAdu (3M, 4F) | |
|---|---|---|---|---|---|---|---|---|---|
| **Analyte** | **Measure** | **Mean** | **SD** | **Mean** | **SD** | **Mean** | **SD** | **Mean** | **SD** |
| **Aβ40** | Baseline (pg/mL) | 131.0 | 34.41 | 185.4 | 46.67 | 147.8 | 50.42 | 122.7 | 47.61 |
| | Terminal (pg/mL) | 140.6 | 31.36 | 186.5 | 53.98 | 211.8 | 25.11 | 182.7 | 79.01 |
| | % Baseline | 116.6 | 47.56 | 108.2 | 51.22 | 163.4 | 68.83 | 177.2 | 117.9 |
| **Aβ42** | Baseline (pg/mL) | 20.49 | 5.655 | 27.49 | 6.701 | 22.43 | 10.79 | 19.96 | 7.701 |
| | Terminal (pg/mL) | 51.46 | 19.08 | 73.93 | 31.05 | 77.4 | 20.12 | 77.54 | 35.86 |
| | % Baseline | 276.0 | 155.9 | 286.9 | 157.6 | 401.4 | 207.6 | 421.9 | 251.7 |
| **Aβ42:40** | Baseline (pg/mL) | 0.1577 | 0.0224 | 0.1498 | 0.0211 | 0.1555 | 0.0374 | 0.1655 | 0.0256 |
| | Terminal (pg/mL) | 0.3568 | 0.0535 | 0.3887 | 0.0583 | 0.4015 | 0.0944 | 0.416 | 0.0906 |
| | % Baseline | 129.1 | 44.17 | 160.2 | 24.48 | 172.3 | 60.4 | 156.9 | 69.99 |

Plasma amyloid-β (Aβ) concentrations at baseline and terminal timepoints in mice receiving IgG or chAdu under weekly or single-dose regimens. Data are presented as mean ± SD. Groups include weekly dosing with 30 mg/kg IgG, 1 mg/kg chAdu, or 30 mg/kg chAdu, and a single-dose 30 mg/kg chAdu cohort (n = 3 males and 4 females per group). Baseline and terminal plasma concentrations are shown for Aβ40 and Aβ42, along with the Aβ42:40 ratio. Percent of baseline was calculated for each analyte within each animal prior to group averaging.

Plasma Aβ concentrations increased from baseline to terminal timepoints across all groups, with more pronounced elevations in chAdu-treated animals compared to IgG controls (Table S5). Increases in both Aβ40 and Aβ42 were observed, accompanied by corresponding increases in the Aβ42:40 ratio. The largest percent changes were observed in the 30 mg/kg chAdu groups, with similar trends following both weekly and single-dose administration.

**Table S6. Pilot study plasma Aβ concentrations in ADA-negative mice only.**

| | | Weekly 30 mg/kg IgG (3M, 4F) | | Weekly 1 mg/kg chAdu (1M, 4F) | | Weekly 30 mg/kg chAdu (3M, 2F) | | Single 30 mg/kg chAdu (3M, 4F) | |
|---|---|---|---|---|---|---|---|---|---|
| **Analyte** | **Measure** | **Mean** | **SD** | **Mean** | **SD** | **Mean** | **SD** | **Mean** | **SD** |
| **Aβ40** | Baseline (pg/mL) | 131.0 | 34.41 | 198.2 | 49.44 | 148.6 | 52.81 | 122.7 | 47.61 |
| | Terminal (pg/mL) | 140.6 | 31.36 | 204.6 | 53.76 | 209.3 | 18.75 | 182.7 | 79.01 |
| | % Baseline | 116.6 | 47.56 | 114.2 | 60.61 | 159.5 | 63.95 | 177.2 | 117.9 |
| **Aβ42** | Baseline (pg/mL) | 20.49 | 5.655 | 30.24 | 5.847 | 22.46 | 8.382 | 19.96 | 7.701 |
| | Terminal (pg/mL) | 51.46 | 19.08 | 81.8 | 33.99 | 85.16 | 18.5 | 77.54 | 35.86 |
| | % Baseline | 276.0 | 155.9 | 296.1 | 190.4 | 379.7 | 128.2 | 421.9 | 251.7 |
| **Aβ42:40** | Baseline (pg/mL) | 0.1577 | 0.0224 | 0.1556 | 0.0216 | 0.1635 | 0.014 | 0.1655 | 0.0256 |
| | Terminal (pg/mL) | 0.3568 | 0.0535 | 0.3909 | 0.0699 | 0.4538 | 0.0302 | 0.416 | 0.0906 |
| | % Baseline | 129.1 | 44.17 | 150.8 | 22.23 | 188.9 | 39.87 | 156.9 | 69.99 |

Plasma amyloid-β (Aβ) concentrations at baseline and terminal timepoints in ADA- mice from the pilot PK/PD study. Data are presented as mean ± SD. Groups include weekly dosing with 30 mg/kg IgG, 1 mg/kg chAdu, or 30 mg/kg chAdu, and a single-dose 30 mg/kg chAdu cohort (group sizes indicated for males and females). Baseline and terminal plasma concentrations are shown for Aβ40 and Aβ42, along with the Aβ42:40 ratio. Percent of baseline was calculated for each animal prior to group averaging. Only animals confirmed to be ADA- were included in these analyses. Corresponding analyses including all animals regardless of ADA status are shown in Table S5.

Restricting analyses to ADA- mice (Table S6) yielded patterns of plasma Aβ changes consistent with those observed in the full cohort (Table S5), with increases in Aβ40, Aβ42, and the Aβ42:40 ratio across chAdu-treated groups. Exclusion of ADA+ animals reduced variability within groups and increased the magnitude of treatment-associated differences, resulting in clearer separation between treatment groups. These findings suggest that ADA status contributes to variability in pharmacokinetic and pharmacodynamic measures but does not alter the overall interpretation of treatment effects.

**Table S7. Pilot study brain Aβ concentrations in all study mice and the ADA-negative subset.**

| | | Weekly 30 mg/kg IgG | | Weekly 1 mg/kg chAdu | | Weekly 30 mg/kg chAdu | | Single 30 mg/kg chAdu | |
|---|---|---|---|---|---|---|---|---|---|
| **Analyte** | **Subset** | **Mean** | **SD** | **Mean** | **SD** | **Mean** | **SD** | **Mean** | **SD** |
| **Soluble Aβ40** | All mice | 0.4237 | 0.1345 | 0.4058 | 0.1157 | 0.5436 | 0.3953 | 0.3474 | 0.1021 |
| | ADA- only | 0.4237 | 0.1345 | 0.4473 | 0.1116 | 0.5748 | 0.4773 | 0.3474 | 0.1021 |
| **Soluble Aβ42** | All mice | 2.138 | 0.8957 | 1.968 | 0.3778 | 1.871 | 0.918 | 1.83 | 0.5637 |
| | ADA- only | 2.138 | 0.8957 | 2.107 | 0.358 | 1.58 | 0.8203 | 1.83 | 0.5637 |
| **Insoluble Aβ40** | All mice | 262.1 | 151.6 | 273.6 | 108.4 | 288.3 | 157.6 | 195.8 | 87.27 |
| | ADA- only | 262.1 | 151.6 | 313.8 | 100.1 | 214 | 112.5 | 195.8 | 87.27 |
| **Insoluble Aβ42** | All mice | 2465 | 1062 | 2623 | 630 | 2474 | 1188 | 1939 | 753.7 |
| | ADA- only | 2465 | 1062 | 2866 | 540.1 | 1907 | 792.1 | 1939 | 753.7 |

Brain amyloid-β (Aβ) concentrations in the pilot PK/PD study, shown for all mice and for the subset of ADA- animals. Data are presented as mean ± SD. Groups include weekly dosing with 30 mg/kg IgG, 1 mg/kg chAdu, or 30 mg/kg chAdu, and a single-dose 30 mg/kg chAdu cohort (group sizes indicated for males and females). Soluble and insoluble Aβ40 and Aβ42 concentrations are shown for each treatment group. Values are reported for the full cohort (“All mice”) and for animals confirmed to be ADA- (“ADA- only”). Corresponding plasma Aβ measures for these cohorts are shown in Tables S5–S6.

Brain Aβ concentrations were similar when comparing all mice to the ADA- subset (Table S7). Overall patterns across treatment groups were preserved, with no consistent directional differences introduced by exclusion of ADA+ animals. However, in select groups, exclusion of ADA+ animals modestly altered mean values and reduced variability, suggesting that ADA status contributes to variability in brain Aβ measures without changing the overall interpretation of treatment effects.
