## Supplementary Methods for "Molecular mechanisms underlying amyloid lowering by aducanumab: differential and comparative effects of sex and IgG reveal the post-treatment disease brain"

Subjects and housing conditions:

Breeding pairs were acquired directly from The Jackson Laboratory (JAX), and subjects enrolled in the studies were the hemizygous 5XFAD offspring and their non-transgenic littermate (WT) controls generated by crossing male 5XFAD mice (congenic on the C57BL6/J background; JAX stock #34848) with female C57BL6/J (JAX stock #000664) mice. Breeders were pair mated at the University of Pittsburgh (Pitt) to generate both the pilot and chronic study cohorts.

5XFAD mice overexpress human APP and five human AD mutations (K670N/M671L, I716V, V717I, M146L and L286V) [1]. This model has been extensively characterized by the IU/JAX/Pitt MODEL-AD Center [2].

Within an AAALAC-certified vivarium, breeding pairs were housed in a designated breeding room. Following weaning, study subjects were group housed by sex ($n$ = 2-4) and transferred to a separate housing room. Subjects were maintained on LabDiet 5P76 in a 12h light/dark cycle with lights on at 7AM. All study procedures were performed during the light cycle. Mouse cages included P.J. Murphy coarse certified Aspen Sani-chip bedding with domes (Bio-serv InnoDome, Cat# S3174) and nestlets (Ancare, Cat# NES3600) for enrichment. Housing rooms were 72-74°F with humidity kept at 40-60%. Ad libitum water was provided through a lixit system at a pH of 4-5 and chlorination levels at 2-3ppm, and water parameters were checked every 2 weeks by animal care staff.

Subjects were excluded from analyses only according to prespecified technical criteria, including incorrect genotype confirmed at terminal collection, detectable drug exposure in vehicle treated controls, or major technical issues (e.g., equipment failure or incorrect treatment administration).

Excluded subjects and rationale are provided in Supplementary Table S1. Pilot and chronic cohort sample sizes are detailed in Supplementary Tables S2 and S3.

Genotyping

Genotyping was confirmed prior to study assignment and upon terminal tissue collection as follows: DNA was extracted from either ear punch (prior to study assignment) or tail snip (terminal) using a lysis buffer containing 25mM NaOH and 0.2mM EDTA. Samples in 75 µL lysis buffer were incubated at 98°C for 1 hour, cooled to room temperature, and then neutralized with 75 µL of a 40mM Tris-HCl solution, pH 5.5. Samples were centrifuged at 4000rpm for 4 minutes at 4°C, and a 1:5 dilution of the resulting supernatant was made using sterile water. Transgenic 5XFAD and non-carrier littermates were identified using PS1 forward (5'-AAT AGA GAA CGG CAG GAG CA-3') and reverse (5'-GCC ATG AGG GCA CTA ATC AT-3') primers and internal control forward (5'-CTA GGC CAC AGA ATT GAA AGA TCT-3') and reverse (5'-GTA GGT GGA AAT TCT AGC ATC ATC C-3') primers. Samples were run on 1% agarose gels with 324 bp internal control fragments amplified for non-carrier littermates and both 324 and 608 bp fragments amplified for 5XFAD carriers.

Detection of chAdu in plasma and brain by mass spectrometry

All raw and processed targeted mass spectrometry data are available on Panorama Web Public and are accessible at the following repositories: https://panoramaweb.org/Indiana%20U%20-%20Proteome%20Analysis/Targeted%20quantitation%20of%20chimeric%20aducanumab%20in%20mouse%20plasma%20and%20cortex/project-begin.view

and

https://panoramaweb.org/Indiana%20U%20-%20Proteome%20Analysis/Reproducability%20and%20repeatability%20Aducanumab%20Assay/project-begin.view

Briefly, stable and light peptides corresponding to sequences in the heavy and light chain of ChAducanumab were synthesized by CPC scientific (Rocklin, CA) at >95% purity for method development. Proteins were extracted from mouse cortex hemispheres following cryohomogenization by sonication in a Bioruptor in 20x volume of 40 mM HEPES, 350 mM NaCl, 0.4% NP-40 with HALT protease inhibitor (Thermo Fisher Scientific). Following 20 min centrifugation at 12,000 rcf, the soluble protein was removed from pellet and 200 µL (considered an equivalent of 10 mg wet weight cortex) transferred to a 96 well plate for the assay described below. Plasma proteins were extracted by mixing 10 µL of plasma with 190 µL of RIPA (Boston Bioproducts, Inc) plus HALT protease inhibitor. Half of this was used for the following assay, or an equivalent of 5 µL plasma.

Protein A, reduction/alkylation, tryptic digestion

The workflow was entirely performed using an Agilent Assaymap Bravo (Agilent) and 96 well plate formats. Following equilibration, 5 µL PAW cartridges A tips were loaded with the equivalent of 10 mg cortex extract or 5 µL plasma extract. Tips were washed first with 10x volume of extraction buffer, then with 10x volume PBS (Thermo Scientific), and then eluted with 5x volume 0.5% trifluoracetic acid (TFA). Samples were neutralized using 1 M Tris, pH 9.0, then denatured and reduced with the addition of 8 M Urea and a final concentration of 50 mM TCEP (Sigma Aldrich). After 30 min at room temperature, a final concentration of 100 mM CAA (Sigma Aldrich) and incubation for 30 min covered at room temperature. The urea was diluted to <2 M and 1 μg trypsin/LysC (Promega) was added before the plate was sealed and incubated overnight at 37 °C with 600 rpm shaking. Samples were then acidified and approximately 1/3 of each cortex digest or 1/15th of each plasma digest were loaded on Evotips with the equivalent of 5 fmol (pilot study) or 10 fmol (PD study) SIL peptide spike.

PRM assay

Targeted mass spectrometry was performed using the 100 SPD method on an Evosep with EV1064 endurance column and butterfly column jacket set to 30 °C and the mass spectrometer parameters described below. A beta method high organic blank was run between each sample to prevent carryover. A Lumos Tribrid Orbitrap (Thermo Fisher) was used to collect data and Skyline software (MacCoss laboratory) was used for data analysis. The mass spectrometer parameters were as follows: MS1 scan 0-10min, APD on, default charge state 2, orbitrap res 120k, quadrupole isolation, range 620-990 m/z, RF lens 30%, AGC target 100%, max IT automatic, profile. tMS2 OTHCD scans: 3-9 min, Quadrupole isolation, 2 m/z window, fixed HCD 30%, orbitrap res 30k, normal mass range, auto scan range, 30% RF, AGC target 200 %, max IT 200 ms, centroided, loop control ALL, Dynamic RT off, time mode start/end:

Compound m/z z t start (min) t stop (min)

LLIY_Lchain_2 831.4647 2 6.3 8.4

LLIY_SIL_Adu_Lchain_2 836.4688 2 6.3 8.4

AED_Adu_Hchain_2 659.7903 2 3.18 5.18

AED_SIL_Adu_Hchain_2 664.7944 2 3.18 5.18

DIQ_Adu_Lchain_1_Ox 947.9442 2 3.68 5.68

DIQ_SIL_Adu_Lchain_1_Ox 952.9483 2 3.68 5.68

DIQ_Adu_Lchain_1_NoOx 939.9467 2 4.76 6.76

DIQ_SIL_Adu_Lchain_1_NoOx 944.9509 2 4.76 6.76

Each plate was run with a standard curve of chAdu spiked into 5XFAD cortex or plasma extract and final chAdu concentrations were calculated using a minimum of 3 process replicate standard curves. Although all peptides tracked in relative abundance, the AED peptide was utilized for the PK/PD determination studies as it had low background, low carryover from high samples, and a minimum of 3 orders of magnitude dynamic linear range in both cortex and plasma enriched extracts.

Endotoxin assay

To verify the absence of endotoxins in chAdu and IgG control vials prepared for chronic dosing, we used Endozyme II kits (bioMerieux, Cat# 890030). Stock antibody vials and diluted formulations were tested in duplicate along with endotoxin standards, positive and negative controls according to kit instructions.

Anti-drug antibody screening

For these studies, positive control sera generated from immunization of New Zealand White rabbits with chAdu were sourced from Pacific Immunology (Ramona, CA). Pre-immunization and post-immunization sera were used as negative and positive controls, respectively, along with drug-naïve 5XFAD mouse plasma negative controls, during ADA assay development and study sample testing. Positive ADA screening results were determined as average (sample signal) > screening cut point (1.16 × mean negative control signal).

All incubations throughout the protocol were performed with moderate shaking at room temperature (RT) unless otherwise noted.

To capture and elute ADAs for screening, clear flat-bottom Maxisorp 96-well plates (Thermo Scientific, Cat# 442404) were coated with 5 μg/mL chAdu diluted in BupH carbonate-bicarbonate buffer (Thermo Scientific, Cat# 28382) incubated at 100 μL per well for 1 hour. After coating, plates were washed 3 times with 1X tris-buffered saline containing 0.05% Tween-20 (TBST). Then, 50 μL of 0.25M Tris-HCl, pH 9, was added to each well. Plasma samples were diluted 1:10 in tris-buffered saline (TBS) and acidified with 300mM acetic acid at a ratio of two parts diluted plasma to one-part acetic acid. Each sample was inverted at RT for 5 minutes to separate ADAs from residual circulating drug. 100 μL of acidified samples were added to each buffered well in the chAdu-coated 96-well plates, and the plates were incubated over night at 4°C, shaking, to capture ADAs. The following day, each plate was washed 3 times with TBST, and 65 μL of 300mM acetic acid was added to each well and incubated for 5 minutes. After elution, 30 μL of 0.25M Tris-HCl, pH9, was added to each well to neutralize each sample.

To create ADA bridging complexes, first, chAdu was biotinylated using EZ-Link Sulfo-NHS-LC-Biotinylation Kit (Thermo Scientific, Cat# 21435), and separately, chAdu was bound to digoxygenin (DIG) using ChromaLINK Digoxigenin On-Shot Antibody Labeling Kit (Vector Laboratories, Cat# B-9014-009K). 225 μL of phosphate-buffered saline (PBS) containing Blocker Casein (Thermo Scientific, Cat# 37582) and a mixture of 1 μg/mL biotinylated chAdu and 1 μg/mL DIG-bound chAdu was added to each well of a fresh 96-well polypropylene plate (Thermo Scientific, Cat#267334), and 25 μL of each eluted sample was added to the plate. The plates were then incubated overnight at 4°C with gentle shaking to create biotin-chAdu, ADA, DIG-chAdu complexes.

After bridging, all samples and controls were run in duplicate on streptavidin-coated Nunc plates (Thermo Scientific, Cat# 15500) to screen for the presence of ADAs. Each plate was washed 3 times with 1X PBS containing 0.05% Tween-20 (PBST), and 100 µL per well of each bridging complex solution was incubated for 30 minutes. The plates were again washed with 1X PBST three times, and 100 µL 1:2000 HRP-conjugated anti-DIG antibody (Immunology Consultants, Catalog No. CDIG65P) in PBST was added to each well and incubated for 30 minutes. After 3 more washes in PBST, 100 µL TMB substrate solution (Thermo Scientific, Cat# N301) was added per well and incubated for 15-30 minutes. When desired color developed, 100 µL TMB Stop solution (Thermo Scientific, Cat# N600) was added to each well and mixed for 30-60 seconds. Absorbance was read at 450 nm on a SpectraMax i3x plate reader.

Pilot behavioral assessment of IgG effects

In parallel with the PK/PD pilot, a separate cohort of 9-month aged male WT littermates was enrolled in a behavioral study to investigate the potential effects of IgG on behavior. For these studies saline (0.9% NaCl) and IgG2a (30 mg/kg) were blinded as treatment groups N and O, respectively, and administered weekly via i.p. injection. Beginning at week 7, behavioral testing was conducted with 1 test per week 24 hours after treatment with IgG or NaCl as follows: Frailty (Week 7), open field (Week 8), rotarod motor coordination (Week 9), and spontaneous alternation spatial working memory task (Week 10).

Pilot PK characterization and post hoc population modeling

An initial 4-week pilot PK study was conducted in 8- to 9-month-old male and female 5XFAD mice and age- and sex-matched non-carrier (WT) littermates (Fig. 1A). The pilot study comprised five arms: chAdu 1 mg/kg once weekly in 5XFAD mice; chAdu 30 mg/kg once weekly in 5XFAD mice; chAdu 30 mg/kg as a single dose followed by saline for the remaining 3 weeks in 5XFAD mice; murine IgG2aκ isotype control 30 mg/kg once weekly in 5XFAD mice; and murine IgG2aκ isotype control 30 mg/kg once weekly in WT littermates (n = 3-4/sex/arm for the chAdu and 5XFAD IgG arms; n = 4-5/sex for the WT IgG arm). All injections were administered intraperitoneally at 10 mL/kg. Baseline plasma (pre-drug treatment) was collected via the tail tip method [3] into 50 μL EDTA-coated capillary tubes (Sarstedt, Cat#17.2113.150) 10 days before initiation of dosing, and Aβ40 and Aβ42 were measured using V-PLEX Aβ Panel 1 (4G8) (Meso Scale Discovery, Cat# K15199E). Baseline plasma Aβ42:40 was used to counterbalance subjects into treatment groups to minimize bias by ensuring the average plasma Aβ42:40 in any one group was not statistically different from any other group. Once counterbalanced, groups were randomized into five blinded treatment groups. Mice were weighed immediately before dosing, and that day's body weight was used to calculate the precise dosage for administration (mg/kg).

For PK analysis, blood was collected via tail tip method into 50 μL heparin coated microhematocrit capillary tubes (Fisherbrand, Cat# 22-362566), immediately placed on ice in labeled and chilled 1.5 mL Eppendorf tubes and then centrifuged 20-60min after collection at 4°C and 14,500 rpm for 10 minutes. After centrifugation, 25 μL plasma was carefully removed and aliquoted in a new, chilled 1.5 mL tube, and the remaining pellet was discarded. After aliquoting, plasma was quickly frozen on dry ice and then stored at -80°C until analysis. Plasma PK sample collection times were as follows: immediately before first dose then subsequently at 24h and 72h after first dose (days 0, 1 and 3), and immediately before and 72h after the second, third and fourth weekly doses (days 7, 10, 14, 17, 21 and 24) (Fig. 1A). Terminal plasma, brain, and tail tissue from the pilot study were collected (as described below in Terminal Tissue Collection) on day 28, 7 days after the final dose which was administered on day 21.

The pilot study included 1 and 30 mg/kg arms to characterize chAdu exposure across the pharmacologic range relevant to chronic dosing [4]. The plasma concentration-time data were used to develop a population pharmacokinetic (popPK) model and perform steady-state exposure simulations for post hoc characterization of chronic-study exposure.

The popPK model was developed in R (version 4.5.2) using nlmixr2 (version 5.0.0) with the first-order conditional estimation with interaction (FOCEI), algorithm in R (version 4.5.2). The model-building dataset comprised the three chAdu pilot arms (30 mg/kg single dose, 1 mg/kg once weekly, and 30 mg/kg once weekly; N = 21 mice; 12 females and 9 males). Five observations were excluded prior to fitting as suspected assay or sampling artifacts (ID 27 at day 24; ID 21 at days 21, 24, and 28; ID 31 at day 7). Four of these observations were more than two orders of magnitude below the corresponding arm mean.

A one-compartment disposition model with first-order absorption and first-order elimination was parameterized in terms of apparent clearance (CL/F) and apparent central volume (V/F). Because the first post-dose sample was collected approximately 1 day after dosing and the observed Tmax was 1 day across subjects, the absorption rate constant (Ka) for intraperitoneal dosing was fixed at 1 $day^{-1}$. Between-subject variability was modeled as exponential random effects on CL/F and V/F, and residual variability was modeled with a proportional error structure after evaluation against a combined additive-and-proportional alternative. Sex and baseline body weight were evaluated as candidate covariates on $\eta CL$ and $\eta V$ using post hoc empirical Bayes estimates from the base model (Wilcoxon rank-sum test for sex, linear regression for body weight); the predefined retention criterion was $p < 0.05$. Model evaluation included standard goodness-of-fit diagnostics, individual concentration-time fits, and visual predictive checks stratified by pilot arm using 1000 simulated replicates of the dataset.

Final-model parameter estimates were used post hoc to simulate once-weekly intraperitoneal dosing at 0.1, 1.56, and 30 mg/kg over 17 weeks for a population of 500 virtual subjects, incorporating between-subject variability on CL/F and V/F from the estimated variance-covariance matrix. Steady-state exposure metrics (Cmax,ss, Ctrough,ss, and AUCτ,ss) were derived from the final simulated dosing interval at the typical-value level. Observed chronic cohort plasma chAdu concentrations collected at weeks 8, 11, and 17 (48 hours post-dose) were overlaid on the simulated 90% prediction intervals to provide a post hoc visual assessment of model predictive performance at chronic steady state (Supplementary Fig. S5E).

<u>Chronic treatment of chAdu and IgG</u>

The chronic treatment period was carried out for 17 weeks (Fig. 1A). Three dose levels were administered once weekly: 0.1 mg/kg as a sub-pharmacologic dose (~30-fold below the 3 mg/kg efficacy benchmark in Tg2576 mice), 1.56 mg/kg as a near-threshold dose (~2-fold below), and 30 mg/kg as a robustly active dose matching the highest previously tested level [10]. Male and female 5XFAD mice were assigned to one of five chronic treatment arms: chAdu 0.1 mg/kg once weekly, chAdu 1.56 mg/kg once weekly, chAdu 30 mg/kg once weekly, IgG2aκ isotype control 1.56 mg/kg once weekly, or saline (n = 10-12/sex/arm). Age- and sex-matched WT littermates received IgG2aκ isotype control 1.56 mg/kg once weekly and served as a control for behavioral assays. The saline-treated 5XFAD arm was included as a negative control to differentiate IgG-mediated effects from chAdu-specific effects on amyloid, transcriptomic, and proteomic outcomes. The study was conducted as two batched cohorts balanced by treatment and sex (n = 5-

6/sex/treatment) staggered 4 weeks apart to allow for appropriate timing of behavioral assessments and terminal tissue collections which were conducted precisely 48 hours after the $17^{th}$ (final) dose. Prior to the first dose, baseline EDTA plasma was collected following the same procedures as described above for the pilot studies and measured for Aβ40 and Aβ42 concentrations. These data were used to counterbalance the treatment groups based on plasma Aβ42:40 ratios, such that the group mean for pre-treatment plasma Aβ42:40 was not significantly different or biased across any treatment group. Subjects were then assigned to blinded treatment groups. Plasma for PK and/or PD measures was collected 48 hours after dosing on weeks 4, 8 and 11. Plasma samples collected with EDTA-coated capillary tubes (Sarstedt, Cat#17.2113.150) on weeks 4 and 8 were used for Aβ quantification, and EDTA plasma samples collected on week 11 were used for immunogenicity screening. Heparinized plasma collected on weeks 8 and 11 were used for PK analysis. At the conclusion of chronic treatment, terminal cerebrospinal fluid (CSF), blood, brain, and tail tissue were collected 48 hours after final dosing and stored at -80°C.

Terminal tissue collection

Mice were anesthetized to the surgical plane with isoflurane and remained anesthetized throughout CSF collection. The cisterna magna was surgically exposed and then punctured with a glass capillary tube. Approximately 5 µL of CSF was collected from each mouse. Any CSF sample contaminated by blood was discarded. Following CSF collection, mice were decapitated, and one 50 µL capillary tube of heparinized plasma was collected from the trunk blood, place in a chilled 1.5 mL tube, and processed for later PK analysis as described above. The remaining trunk blood was decanted into chilled 500 µL EDTA-coated BD Microtainer™ collection tubes (BD, SKU: 365974) and kept on ice until centrifugation at 4°C for 20min at 14,500 rpm. After centrifugation, plasma was carefully aliquoted for Aβ quantification, immunogenicity screening, and multi-omic analyses, and aliquots were quickly frozen on dry ice and stored at -80°C. After blood collection, brains were dissected from the skull and briefly rinsed (<30 seconds) in ice cold 1X PBS. After rinsing, the brain was sectioned as described below, then frozen and stored at -80°C for later PK, PD, and proteomic assays. To confirm genotypes, 1-2mm of tail tissue was collected at terminal.

#### Tissue processing for multi-omic analyses

During terminal tissue collection, the right hemisphere (minus the cerebellum) from each subject was cross chopped using razor blades on an ice-cold metal platform. The chopped brain was mixed, divided into three even aliquots, transferred to three separate chilled 1.5 mL tubes, and quickly frozen on dry ice, then stored at -80°C. Frozen samples were shipped to Emory University for proteomic analysis and to JAX for transcriptomic analysis.

#### Tissue processing for soluble and insoluble brain fractionation and PK analysis

During terminal collection, the left cerebrum (hemi-brain) was separated from the cerebellum, transferred to a chilled 1.5 mL tube, and quickly frozen on dry ice, then stored at -80°C until processing. Each frozen hemi-brain was transferred to a TT1 tissue TUBE (Covaris, PN520001), submerged in liquid nitrogen for 1 min, then placed into the center of a CP02 cryoPREP automatic dry pulverizer (Covaris) and pulverized while frozen. The TT1 tubes were returned to liquid nitrogen, and each pulverized hemi-brain was mixed to distribute the powder and create two equal aliquots of the mixed brain powder in new 1.5 mL tubes, chilled on dry ice. Samples were stored at -80°C until analysis.

Brain and plasma analysis of Aβ42 and Aβ40 by multi-plex ELISA

For soluble and insoluble protein fraction preparation, we adapted methods previously described in Casali et al, 2017 [3,5]. One aliquot of cryoPREP pulverized hemi-brain was weighed on a scale tared to the weight of an empty, labeled 1.5 mL tube. 1 mL of cold tissue homogenization buffer (THB) containing protease and phosphatase inhibitors was added to the pulverized brain per 100 mg tissue weight, up to a maximum volume of 850 µL, and the sample was mechanically homogenized using a 1.5 mL tube pellet mixer. Soluble and insoluble fractions were subsequently prepared by diethanolamine (DEA) and formic acid (FA) extraction, respectively. Aβ40 and Aβ42 were analyzed in brain fractions and plasma using the Meso Scale Discovery V-PLEX Aβ Peptide Panel 1 (4G8) #K15199E. For analysis of brain fractions, total protein was measured using Pierce Detergent Compatible Bradford Assay Kit (ThermoScientific, Cat# 23246) within brain soluble and insoluble fractions run in duplicate. To dilute buffer components to within the compatible range for the Bradford reagent, we ran soluble fractions at a 2x dilution in sterile water according to the Standard Microplate Protocol with a working range of 100-1500 µg/mL, and we ran insoluble fractions at a 20x dilution in sterile water according to the Micro Microplate Protocol with a working range of 2-25 µg/mL. Soluble and insoluble fraction Aβ concentrations were normalized to their respective total protein for each sample.

Behavioral testing

Following 12 weeks of treatment in the chronic dosing study, behavioral tests were conducted at PITT as previously reported [3] in the following order: frailty, open field test, spontaneous alternation, rotarod. Dosing was performed on Mondays or Tuesdays with behavior assessments occurring on Wednesday-Friday, allowing at minimum a 2-day rest period between i.p. administration and behavior tests.

Frailty assessment

The frailty assessment was conducted as previously described [2,3]. Following acclimation, subjects were individually evaluated for the absence or presence of 26 characteristic traits and reflexes and scored a 0, 0.5, or 1 (based on presence/absence, and severity) for each assessment by a trained observer, blind to genotype and treatment. The frailty index score was calculated as the cumulative score of all measures with a maximum score of 26. Core body temperature was recorded just prior to the conclusion of the frailty assessment via a glycerol lubricated thermistor rectal probe (Braintree Scientific product# RET 3). Temperature was recorded to the nearest 0.1°C (Braintree Scientific product #TH5 Thermalert digital thermometer).

Open field test

Versamax Open Field Arenas (40 cm x 40 cm x 40 cm; Omnitech Electronics, OH USA) were housed within sound-attenuated chambers with lighting in the testing room and arenas consistent with that of the housing room (~500 lux). Mice were placed individually into the center of the arena and infrared beams recorded distance traveled (cm), vertical activity, and perimeter/center time. Data were collected in 5 min time bins for a duration of 60 min.

Spontaneous alternation

Mice were acclimated to the testing room under ambient lighting conditions (20-50 lux). A clear polycarbonate y-maze (in-house fabricated; arm dimensions 33.65 cm length, 6 cm width, 15 cm height) placed on top of an infrared reflecting background (Noldus, The Netherlands) surrounded by a black floor-to-ceiling curtain to minimize extramaze visual cues was used for this test. Mice were placed in the middle of the start arm (A) facing the center of the y-maze for an 8-min test period and the sequence of entries into each arm was recorded via a ceiling-mounted infrared camera integrated with behavioral tracking software (Noldus Ethovision XT). Percent spontaneous alternation is calculated as the number of triads (entries into each of the three different arms of the maze in a sequence of three without returning to a previously visited arm) relative to the number of alteration opportunities.

Rotarod test for motor coordination

An accelerating Rotarod (Ugo-Basile; model 47600) was used for this test. The lighting in the testing room was consistent with that of the housing room (~ 500 lux). Mice were placed on the rotating rod (4 rpm), which accelerates up to 40 rpm over the course of 300 sec. Each mouse was subjected to 3 consecutive trials with a ~1 min inter-trial interval to allow cleaning of the rod between trials. Latency to fall (sec) was measured. Subjects that fell upon initial placement on the rod, before acceleration began, were scored as 0 sec for that trial.

Data processing and statistical analysis

All data were initially analyzed under coded genotypes during data processing and QC, including maintaining the blind for proteomic and transcriptomic analyses as described below. Prior to unblinding, samples meeting predefined *a priori* exclusion criteria were excluded independent of any statistical or mathematical determination. Specific exclusions included genotyping errors confirmed at terminal tissue collection (e.g., WT animals erroneously assigned as 5XFAD and drug treated) and vehicle-treated controls with detectable levels of chAdu based on PK analysis; these samples were excluded from subsequent PD and multi-omic analyses (see Supplementary Table S1). Primary analyses were conducted in the subset of animals that were ADA-, given the impact of ADAs on systemic drug exposure. Analyses including the full cohort irrespective of ADA status are provided in Supplementary Figures S5-S7 and Supplementary Tables S5-S7.

Statistical analysis for brain and plasma chAdu and Aβ, as well as behavioral assays, was conducted using GraphPad Prism (version 10). Pharmacokinetic (PK) measures were analyzed using ordinary or repeated-measures two-way ANOVA, as appropriate, with Šídák's multiple comparison test to assess sex differences within each dose. To evaluate the impact of immunogenicity on drug exposure, plasma exposure was expressed within each dose group as a percentage of the ADA- mean (set to 100%). ADA-positive (ADA+) and ADA- animals were compared using a two-tailed Mann–Whitney test. Pharmacodynamic (PD) measures (brain and plasma Aβ40 and Aβ42) were analyzed within sex using one-way ANOVA with Dunnett's multiple comparisons test (brain) or two-way repeated-measures ANOVA with Tukey's multiple comparisons test (plasma), comparing antibody-treated 5XFAD groups (IgG and chAdu) to saline-treated 5XFAD controls. Behavioral outcomes were analyzed within sex using one-way or two-way ANOVA as appropriate, with IgG-treated WT mice serving as the reference group for comparisons to IgG- and chAdu-treated 5XFAD mice. Statistical analysis for transcriptomic and proteomic datasets is described in detail below. The blind was revealed only after completion of initial analyses for appropriate statistical interpretation.

<u>RNA isolation, preparation, and sequencing</u>

An aliquot of fresh cross-chopped, frozen hemi-brain and matching EDTA plasma was selected from n = 6/sex/treatment for transcriptomic analysis. Samples were stored at -80°C until RNA isolation, preparation, and analysis. Total RNA was isolated from frozen tissue using the NucleoMag RNA Kit (Macherey-Nagel) and the KingFisher Flex purification system (ThermoFisher). Tissues were homogenized in MR1 buffer (Macherey-Nagel) using a Bead Ruptor Elite (Omni International). RNA isolation was performed according to the manufacturer's protocol. RNA concentration and quality were assessed using the Nanodrop 8000 spectrophotometer (Thermo Scientific) and the RNA ScreenTape Assay (Agilent Technologies). Stranded libraries were constructed using the KAPA mRNA HyperPrep Kit (Roche Sequencing and Life Science), according to the manufacturer's protocol. Briefly, the protocol entails isolation of polyA containing mRNA using oligo-dT magnetic beads, RNA fragmentation, first and second strand cDNA synthesis, ligation of Illumina-specific adapters containing a unique barcode sequence for each library, and PCR amplification. The quality and concentration of the libraries were assessed using the D5000 ScreenTape (Agilent Technologies) and Qubit dsDNA HS Assay (ThermoFisher), respectively, according to the manufacturers' instructions. Libraries were pooled and sequenced by the Genome Technologies core facility at The Jackson Laboratory. All samples were sequenced 150 bp paired-end on an Illumina NovaSeq X Plus using the 10B Reagent Kit (Illumina), targeting 30 million read pairs per sample. Once the data was received the samples were concatenated to have a single file for paired-end analysis.

<u>RNA-Sequencing data processing</u>

RNA-Seq data were processed using nf-core/rnaseq pipeline [https://doi.org/10.5281/zenodo.1400710]. To quantify human *APP* and *PSEN1* transgene expression, we created a chimeric mouse genome by concatenating human APP (Chromosome 21: 25880550–26171128; build GRCh38.p10) and PSEN1 (Chromosome 14: 73136418–73223691; build GRCh38.p10) gene sequences into the mouse genome (Ensembl Genome Reference Consortium, build 38) as separate chromosomes (labeled chromosomes 21 and 22). Subsequently, we added gene annotations for human *APP* and human *PSEN1* genes in the resulting gtf file. A STAR index was built from this mouse chimeric genome sequence for alignment. Finally, reads were mapped to the chimeric mouse genome using STAR [6] and gene expression was quantified with RSEM [7]

Proteomic analysis

A separate frozen cross-chopped hemi-brain aliquot from the same subjects (n=6/sex/treatment) used for transcriptomic analysis was shipped to Emory University for proteomic analysis and matched within subject to CSF and plasma. Frozen samples were homogenized in 8 M urea lysis buffer (8 M urea, 10 mM Tris, 100 mM $NaH_2PO_4$, pH 8.5) with HALT protease and phosphatase inhibitor cocktail (ThermoFisher) using a Bullet Blender (NextAdvance) as described [8]. Each Rino sample tube (NextAdvance) was supplemented with ~100 µL of stainless-steel beads (0.9 to 2.0 mm blend, NextAdvance) and 300 µL of lysis buffer. Tissues were added immediately after excision and homogenized with bullet blender at 4 °C with 2 full 5 min cycles. The lysates were transferred to new Eppendorf Lobind tubes and sonicated for 3 cycles consisting of 5 s of active sonication at 30% amplitude, followed by 15 s on ice. Samples were then centrifuged for 5 min at 15,000 x g and the supernatant transferred to a new tube. Protein concentration was determined by bicinchoninic acid (BCA) assay (Pierce). For protein digestion, 100 µg of each sample was aliquoted and volumes normalized with additional lysis buffer. Samples were reduced with 5 mM dithiothreitol (DTT) at room temperature for 30 min, followed by 10 mM iodoacetamide (IAA) alkylation in the dark for another 30 min. Lysyl endopeptidase (Wako) at 1:25 (w/w) was added, and digestion allowed to proceed overnight. Samples were then diluted 7-fold with 50 mM ammonium bicarbonate. Trypsin (Promega) was then added at 1:25 (w/w) and digestion proceeded overnight. The peptide solutions were acidified to a final concentration of 1% (vol/vol) formic acid (FA) and 0.1% (vol/vol) trifluoroacetic acid (TFA) and desalted with a 30 mg HLB column (Oasis). Each HLB column was first rinsed with 1 mL of methanol, washed with 1 mL 50% (vol/vol) acetonitrile (ACN), and equilibrated with 2×1 mL 0.1% (vol/vol) TFA. The samples were then loaded onto the column and washed with 2×1 mL 0.1% (vol/vol) TFA. Elution was performed with 2 volumes of 0.5 mL 50% (vol/vol) ACN.

<u>Isobaric tandem mass tag (TMT) peptide labeling</u>

Each sample (containing 100 μg of peptides) was re-suspended in 100 mM TEAB buffer (100 μL). The TMT labeling reagents (5 mg) were equilibrated to room temperature, and anhydrous ACN (256 μL) was added to each reagent channel. Each channel was gently vortexed for 5 min, and then 41 μL from each TMT channel was transferred to the peptide solutions and allowed to incubate for 1 h at room temperature. The reaction was quenched with 5% (vol/vol) hydroxylamine (8 μL) (Pierce). All channels were then combined and dried by SpeedVac (LabConco) to approximately 150 μL and diluted with 1 mL of 0.1% (vol/vol) TFA, then acidified to a final concentration of 1% (vol/vol) FA and 0.1% (vol/vol) TFA. Labeled peptides were desalted with a 200 mg C18 Sep-Pak column (Waters). Each Sep-Pak column was activated with 3 mL of methanol, washed with 3 mL of 50% (vol/vol) ACN, and equilibrated with 2×3 mL of 0.1% TFA. The samples were then loaded, and each column was washed with 2×3 mL 0.1% (vol/vol) TFA, followed by 2 mL of 1% (vol/vol) FA. Elution was performed with 2 volumes of 1.5 mL 50% (vol/vol) ACN. The eluates were then dried to completeness using a SpeedVac. High pH off line fractionation was conducted as described[9] Dried samples were re-suspended in high pH loading buffer (0.07% vol/vol NH4OH, 0.045% vol/vol FA, 2% vol/vol ACN) and loaded onto a Water's BEH 1.7 um 2.1mm by 150mm. An Thermo Vanquish was used to carry out the fractionation. Solvent A consisted of 0.0175% (vol/vol) NH4OH, 0.01125% (vol/vol) FA, and 2% (vol/vol) ACN; solvent B consisted of 0.0175% (vol/vol) NH4OH, 0.01125% (vol/vol) FA, and 90% (vol/vol) ACN. The sample elution was performed over a 25 min gradient with a flow rate of 0.6 mL/min. A total of 192 individual equal volume fractions were collected across the gradient and subsequently pooled by concatenation into 96 fractions and dried to completeness using a SpeedVac.

LC-MS/MS methods

All fractions were resuspended in an equal volume of loading buffer (0.1% FA, 0.03% TFA, 1% ACN) and analyzed by liquid chromatography coupled to tandem mass spectrometry. Peptide eluents were separated on a custom in-house packed CSH 1.7um (15 cm × 150 μM internal diameter (ID) by a Dionex RSLCnano UPLC (ThermoFisher Scientific). Buffer A was water with 0.1% (vol/vol) formic acid, and buffer B was 80% (vol/vol) acetonitrile in water with 0.1% (vol/vol) formic acid. Elution was performed over a 32 min gradient with flow rate at 1000 nL/min. The gradient was from 1% to 99% solvent B. Peptides were monitored on an Orbitrap Eclipse mass spectrometer with a high-field asymmetric waveform ion mobility spectrometry (FAIMS Pro) ion mobility source (ThermoFisher Scientific). Two compensation voltages (CV) were chosen for the FAIMS. For each CV (-45 and -65) top speed cycle of 1.5 seconds, the full scan (MS1) was performed with an m/z range of 410-1600 at 60,000 resolution at standard settings. The higher energy collision-induced dissociation (HCD) tandem scans were collected at 35% collision energy with an isolation of 0.7 m/z, a resolution of 30,000 with TurboTMT on, an AGC setting of 250% normalized agc target, and a maximum injection time of 54 ms. Dynamic exclusion was set to exclude previously sequenced peaks for 15 seconds within a 10-ppm isolation window.

Datasets (672 raw files (n=7 batches of 96 high pH fractions)) were searched using FragPipe (version 20.0). The FragPipe pipeline relies on MSFragger (version 3.8) [10,11] or peptide identification and Philosopher (version 5.0.0 [12]) for FDR filtering and downstream processing. The mouse protein database used contains canonical isoforms from Uniprot/Swissprot as of 02/2023. The workflow used in FragPipe followed default TMT-16 plex parameters, used for both TMT-16 and TMT-18 experimental design. Briefly, precursor mass tolerance was -20 to 20 ppm, fragment mass tolerance of 20 ppm, mass calibration and parameter optimization were selected, and isotope error was set to -1/0/1/2/3. Enzyme specificity was set to strict-trypsin and up to two missing trypsin cleavages were allowed. Peptide length was allowed to range from 7 to 50 and peptide mass from either 200 to 5,000 Da. Variable modifications that were allowed in our search included: oxidation on methionine, N-terminal acetylation on protein, and N-terminal acetylation on peptide, with a maximum of 3 variable modifications per peptide. Peptide Spectral Matches were validated using Percolator [13].The false discovery rate (FDR) threshold was set to 1% and protein and peptide abundances were quantified using Philosopher for downstream analysis.

Protein quantitation and normalization

The protein abundances were normalized by scaling total protein signal within each channel for each specific case sample to the maximum channel-specific total signal. We then used a tunable median polish approach, TAMPOR, to remove technical batch variance in the proteomic data, as previously described [14]. TAMPOR is utilized to remove intra-batch and inter-batch variance while preserving meaningful biological variance in protein abundance values, normalizing to the median of selected intra-batch samples. This approach is robust to outliers and columns with up to 50% values missing. If a protein had more than 50% samples with missing values, it was removed from the matrix. No imputation of missing values was performed for any cohort. For the current data, TAMPOR leverages the median protein abundance from the pooled Global Internal Standard (GIS) TMT channels as the denominators in both factors to normalize sample-specific protein abundances across the 5 batches of samples.

Differential gene and protein expression analysis

Differentially expressed genes were identified using the R Bioconductor package DESeq2 (v1.16.1)[15]. We used the Benjamini-Hochberg corrected p-values with a significance threshold of 0.05 to identify differentially expressed genes. Differentially expressed proteins were identified using one-way ANOVA followed by post hoc correction using Tukey HSD test to match methods used in analogous human studies [16].

Generalized linear model

We applied a generalized linear model (GLM) to quantify the effects of sex, genotype, and chAdu dose. These factors were evaluated by fitting the following linear model:

$$log(expr) = \beta_0 + \sum_i \beta_i + \varepsilon$$

The sum is over sex (male), genotype (5XFAD), IgG treatment, and chAdu dose (low, middle, and high) used in this study. The expr represents gene expression measured by RNA-Seq transcript per million (TPM). In this model, IgG-treated WT littermates served as the control group for the

IgG-treated 5XFAD mice, while the saline-treated 5XFAD mice served as controls for the IgG-treated and chAdu-treated 5XFAD groups to estimate the effects of genotype and chAdu dose.

AD-related protein co-expression modules from human postmortem brain tissue

Human protein co-expression modules were obtained from previously published protein expression profiles from human dorsolateral prefrontal frontal cortex [16]. Briefly, Johnson et al. analyzed over 500 dorsal prefrontal cortex (DLPFC) tissues spanning control, asymptomatic AD (AsymAD), and AD cases to construct a comprehensive TMT-based Alzheimer's disease proteomic network using WGCNA. This network consists of 44 protein co-expression modules that capture coordinated expression patterns across healthy and diseased tissue. Johnson, et al. functionally annotated these modules using Gene Ontology analysis of its constituent proteins and assigned cell types to each module, and measured correlations of each summary module eigenprotein to neuropathological or cognitive traits present in the cohorts [16]. Twelve modules were noted to correlate more strongly to AD traits than the others. We obtained sex-adjusted case–control log2 fold-change values for all quantified proteins, along with their corresponding protein co-expression module assignments, from the AD Knowledge Portal (https://www.synapse.org/#!Synapse:syn25453861). The log2 fold-change values for proteins within these modules were originally computed from sex-regressed protein abundance matrices. Therefore, we re-processed the protein abundance matrices to preserve the sex effect by re-running the non-parametric bootstrap regression described by Johnson et al., restricting the adjustment to age at death and postmortem interval (PMI) [16]. This approach ensured that sex-related variation was retained in the data. Then, we calculated case–control log2 fold-change values for all quantified proteins separately for males and females. Finally, we replaced the original sex-regressed log2 fold-change values within each module with sex-specific log2 fold-change values and generated distinct male-specific and female-specific datasets for the protein co-expression modules.

Human AMP-AD gene co-expression modules

Wan et al. (2020) discovered 30 human brain co-expression modules based on the meta-analysis of differential gene expression from seven distinct regions in postmortem samples obtained from three independent LOAD cohorts [9]. These modules were grouped into five distinct consensus clusters representing shared AD-related changes across studies and brain regions [9]. Reactome pathway enrichment analysis was applied to annotate each cluster with distinct biological themes, ranking pathways by Bonferroni corrected *p*-values to ensure statistical rigor. Data for the 30 human brain co-expression modules generated by the Accelerating Medicines Partnership for Alzheimer's Disease (AMP-AD) studies were obtained from the Synapse data repository (https://www.synapse.org/#!Synapse:syn11932957/tables/;). The log2 fold-change values for genes within these modules were originally computed from sex-regressed residualized counts. To enable direct comparisons between gene expression changes in female mice and female AD cases, and between male mice and male AD cases, we replaced these sex-regressed log2 fold-change values with sex-specific log2 fold-change values within each gene module. We obtained case–control log2 fold-change values for all quantified genes separately for males and females from the Synapse data repository (https://www.synapse.org/Synapse:syn30821563) and generated distinct male-specific and female-specific datasets for the 30 human brain co-expression modules.

Mouse-human correlation analysis

We assessed the similarity between mouse and human disease-related expression changes by computing Pearson correlations between log2 fold change values in human AD cases versus controls and the corresponding log2 fold change values in mouse models (saline-treated 5XFAD vs. IgG-treated WT littermates, and chAdu-treated or IgG-treated 5XFAD vs. saline-treated 5XFAD), stratified by sex. Correlations were calculated across the set of orthologous genes in each AMP-AD module [17,18], and across orthologous proteins within corresponding proteomics modules using cor.test function in R as:

$$cor.test\ (log_2FC\ (AD/control),\ log_2FC(model/control))$$

where logFC(AD/control) is the log fold change in transcript or protein expression of human AD patients compared to control patients. We plotted the correlation results using the ggplot2 package in R. Framed circles were used to denote significant ($p < 0.05$) positive (blue) and negative (red) Pearson's correlation coefficients. The color intensity and size of the circles were sized proportional to Pearson's correlation coefficient.

Functional enrichment analysis

Functional annotations and enrichment analyses were performed using the R Bioconductor package clusterProfiler [19], applying the enrichGO function with the org.Mm.eg.db annotation database to evaluate Gene Ontology terms. The significance threshold for all enrichment analyses was set at 0.05 using Benjamini–Hochberg adjusted *p*-values. Gene functional enrichment analyses are informative but sometimes it is difficult to understand how the enriched terms relate to the biology of AD. Cary et al. developed 19 biological domains that capture the AD-associated endophenotypes and defined them using an exhaustive set of Gene Ontology (GO) terms, with the intent to keep each domain siloed in a biologically coherent fashion [20]. The GO enrichment results were organized into biological domains based on the GO identifiers of enriched terms [20].

Correlation with AD biological domains

We grouped genes and proteins into AD-associated biological domain based on their annotation to any Gene Ontology term associated with each biological domain [20]. Next, we assigned the previously calculated log2 fold-change values to each protein within its respective biological domain, separately for males and females. For each biological domain, we then computed Pearson correlations between human AD log2 fold change values and the corresponding log2 fold change values in mouse models, stratified by sex. In transcriptomics datasets, For each biological domain, the Pearson correlation was computed between the estimated effect of the genotype (5XFAD), IgG treatment, and chAdu dose (low, middle, and high) on each gene and the meta-analysis treatment effect (*https://www.synapse.org/#!Synapse:syn22758536/tables/*) for each orthologous genes derived from all human AMP-AD transcriptomic datasets.

### SUPPLEMENTARY REFERENCES:


[1] Oakley H, Cole SL, Logan S, Maus E, Shao P, Craft J, et al. Intraneuronal β-amyloid aggregates, neurodegeneration, and neuron loss in transgenic mice with five familial Alzheimer's disease mutations: Potential factors in amyloid plaque formation. Journal of Neuroscience 2006;26:10129–40. https://doi.org/10.1523/JNEUROSCI.1202-06.2006.

[2] Oblak AL, Lin PB, Kotredes KP, Pandey RS, Garceau D, Williams HM, et al. Comprehensive Evaluation of the 5XFAD Mouse Model for Preclinical Testing Applications: A MODEL-AD Study. Front Aging Neurosci 2021;13. https://doi.org/10.3389/fnagi.2021.713726.

[3] Oblak AL, Cope ZA, Quinney SK, Pandey RS, Biesdorf C, Masters AR, et al. Prophylactic evaluation of verubecestat on disease- and symptom-modifying effects in 5XFAD mice. Alzheimer's and Dementia: Translational Research and Clinical Interventions 2022;8. https://doi.org/10.1002/trc2.12317.

[4] Sevigny J, Chiao P, Bussière T, Weinreb PH, Williams L, Maier M, et al. The antibody aducanumab reduces Aβ plaques in Alzheimer's disease. Nature 2016;537:50–6. https://doi.org/10.1038/nature19323.

[5] Casali B, Landreth G. Aβ Extraction from Murine Brain Homogenates. Bio Protoc 2016;6. https://doi.org/10.21769/bioprotoc.1787.

[6] Dobin A, Davis CA, Schlesinger F, Drenkow J, Zaleski C, Jha S, et al. STAR: ultrafast universal RNA-seq aligner. Bioinformatics 2013;29:15–21. https://doi.org/10.1093/bioinformatics/bts635.

[7] Li B, Dewey CN. RSEM: accurate transcript quantification from RNA-Seq data with or without a reference genome. BMC Bioinformatics 2011;12:323. https://doi.org/10.1186/1471-2105-12-323.

[8] Ping L, Kundinger SR, Duong DM, Yin L, Gearing M, Lah JJ, et al. Global quantitative analysis of the human brain proteome and phosphoproteome in Alzheimer's disease. Sci Data 2020;7:315. https://doi.org/10.1038/s41597-020-00650-8.

[9] Wan Y-W, Al-Ouran R, Mangleburg CG, Perumal TM, Lee T V., Allison K, et al. Meta-Analysis of the Alzheimer's Disease Human Brain Transcriptome and Functional Dissection in Mouse Models. Cell Rep 2020;32:107908. https://doi.org/10.1016/j.celrep.2020.107908.

[10] Kong AT, Leprevost F V, Avtonomov DM, Mellacheruvu D, Nesvizhskii AI. MSFragger: ultrafast and comprehensive peptide identification in mass spectrometry–based proteomics. Nat Methods 2017;14:513–20. https://doi.org/10.1038/nmeth.4256.

[11] Yu F, Teo GC, Kong AT, Haynes SE, Avtonomov DM, Geiszler DJ, et al. Identification of modified peptides using localization-aware open search. Nat Commun 2020;11:4065. https://doi.org/10.1038/s41467-020-17921-y.

[12] da Veiga Leprevost F, Haynes SE, Avtonomov DM, Chang H-Y, Shanmugam AK, Mellacheruvu D, et al. Philosopher: a versatile toolkit for shotgun proteomics data analysis. Nat Methods 2020;17:869–70. https://doi.org/10.1038/s41592-020-0912-y.

[13] Käll L, Canterbury JD, Weston J, Noble WS, MacCoss MJ. Semi-supervised learning for peptide identification from shotgun proteomics datasets. Nat Methods 2007;4:923–5. https://doi.org/10.1038/nmeth1113.

[14] Dammer EB, Seyfried NT, Johnson ECB. Batch correction and harmonization of –Omics datasets with a tunable median polish of ratio. Frontiers in Systems Biology 2023;3. https://doi.org/10.3389/fsysb.2023.1092341.
[15] Love MI, Huber W, Anders S. Moderated estimation of fold change and dispersion for RNA-seq data with DESeq2. Genome Biol 2014;15:550. https://doi.org/10.1186/s13059-014-0550-8.
[16] Johnson ECB, Carter EK, Dammer EB, Duong DM, Gerasimov ES, Liu Y, et al. Large-scale deep multi-layer analysis of Alzheimer’s disease brain reveals strong proteomic disease-related changes not observed at the RNA level. Nat Neurosci 2022;25:213–25. https://doi.org/10.1038/s41593-021-00999-y.
[17] Pandey RS, Graham L, Uyar A, Preuss C, Howell GR, Carter GW. Genetic perturbations of disease risk genes in mice capture transcriptomic signatures of late-onset Alzheimer’s disease. Mol Neurodegener 2019;14:50. https://doi.org/10.1186/s13024-019-0351-3.
[18] Preuss C, Pandey R, Piazza E, Fine A, Uyar A, Perumal T, et al. A novel systems biology approach to evaluate mouse models of late-onset Alzheimer’s disease. Mol Neurodegener 2020;15:67. https://doi.org/10.1186/s13024-020-00412-5.
[19] Yu G, Wang L-G, Han Y, He Q-Y. clusterProfiler: an R Package for Comparing Biological Themes Among Gene Clusters. OMICS 2012;16:284–7. https://doi.org/10.1089/omi.2011.0118.
[20] Cary GA, Wiley JC, Gockley J, Keegan S, Amirtha Ganesh SS, Heath L, et al. Genetic and multi-omic risk assessment of Alzheimer’s disease implicates core associated biological domains. Alzheimer’s & Dementia: Translational Research & Clinical Interventions 2024;10. https://doi.org/10.1002/trc2.12461.
