## Supplementary Figures for "Molecular mechanisms underlying amyloid lowering by aducanumab: differential and comparative effects of sex and IgG reveal the post-treatment disease brain"

**LIST OF SUPPLEMENTARY FIGURES**


| **Figure** | **Description** |
|---|---|
| Figure S1 | Final model goodness-of-fit diagnostics |
| Figure S2 | Individual concentration-time fits for the 21 chAdu-treated 5XFAD mice in the population PK model-building dataset |
| Figure S3 | Empirical Bayes estimates versus screened covariates (sex and baseline body weight) |
| Figure S4 | Visual predictive check stratified by pilot arm |
| Figure S5 | Pharmacokinetic analyses including all mice regardless of ADA status. |
| Figure S6 | Chronic plasma Aβ in the full chronic cohort, including ADA+ animals |
| Figure S7 | Terminal brain Aβ40 and Aβ42 in the full chronic cohort, including ADA+ animals, and absolute brain Aβ concentrations in the ADA- subset |
| Figure S8 | Reciprocal regulation of disease-associated proteins by chAdu and IgG treatment, and sex-correlation of proteomic responses across treatment arms |

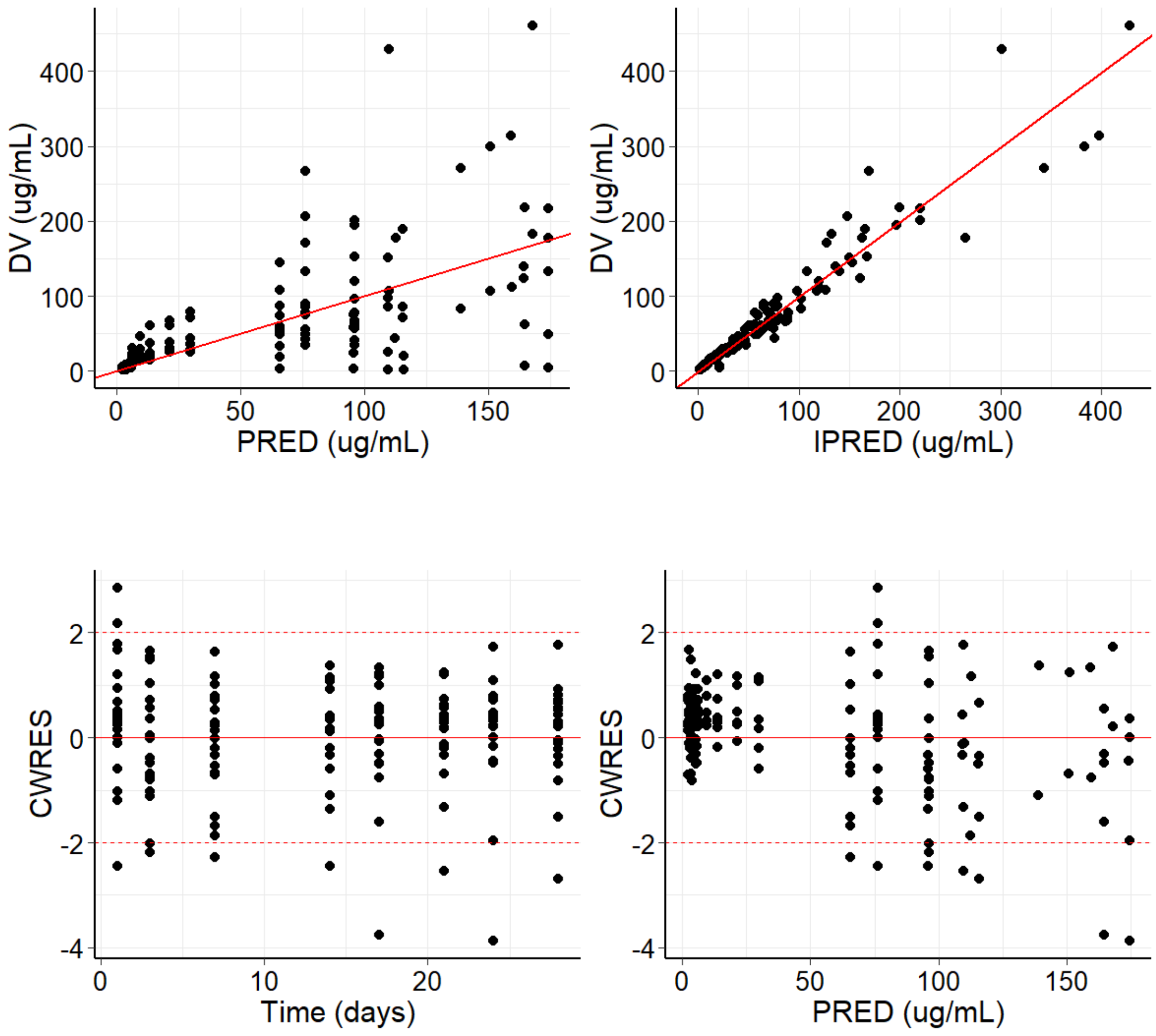


**Figure S1. Final model goodness-of-fit diagnostics.**

Top left, observed plasma chAdu concentration (DV) versus population-predicted concentration (PRED). Top right, DV versus individual-predicted concentration (IPRED). Bottom left, conditional weighted residuals (CWRES) versus time. Bottom right, CWRES versus PRED. Solid red lines indicate the line of unity (top) or zero (bottom); dashed lines mark CWRES = ±2.

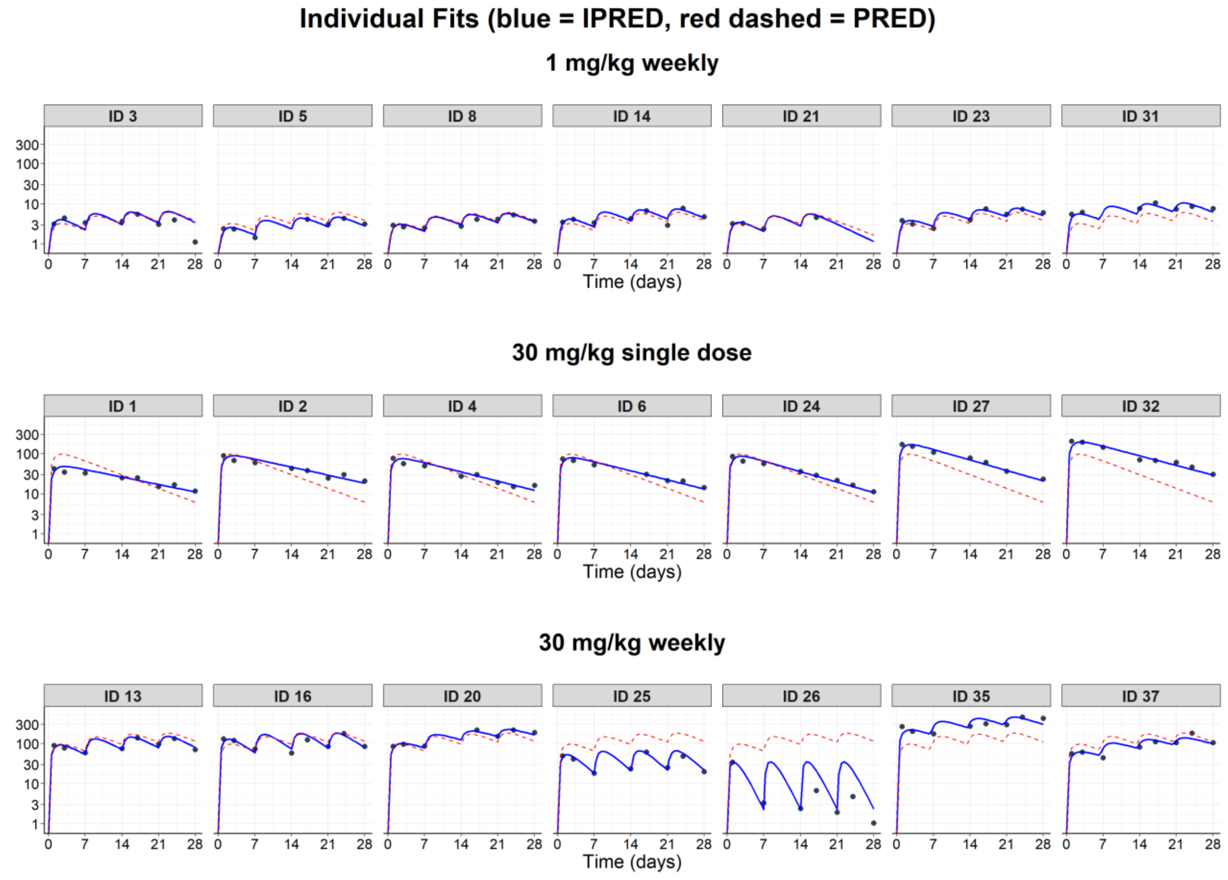


**Figure S2. Individual concentration-time fits for the 21 chAdu-treated 5XFAD mice in the population PK model-building dataset.**

Symbols are observed plasma chAdu concentrations. Solid lines are individual model predictions (IPRED); dashed lines are population predictions (PRED). Panel labels indicate subject ID, sex, dose level, and dosing regimen. Concentration is shown on a $\log_{10}$ scale.

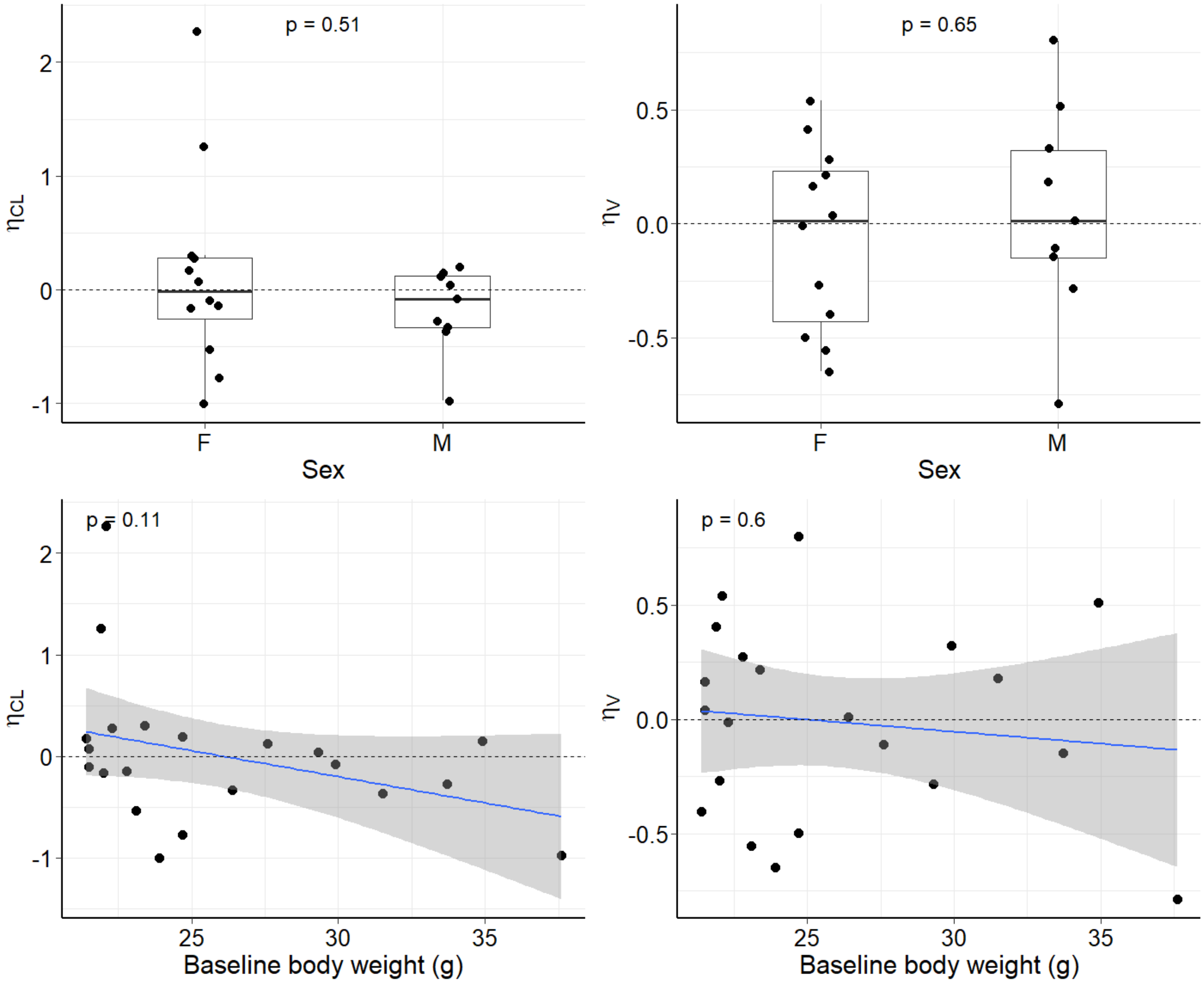


**Figure S3. Empirical Bayes estimates ($\eta_{CL}$, $\eta_V$) versus screened covariates.**

Top, η versus sex (F vs M). Bottom, η versus baseline body weight, with linear regression fit and 95% confidence band. Reported p-values are from Wilcoxon rank-sum tests (sex) or linear regression (body weight). No covariate met the predefined retention criterion ($p < 0.05$); the base model was retained as the final model.

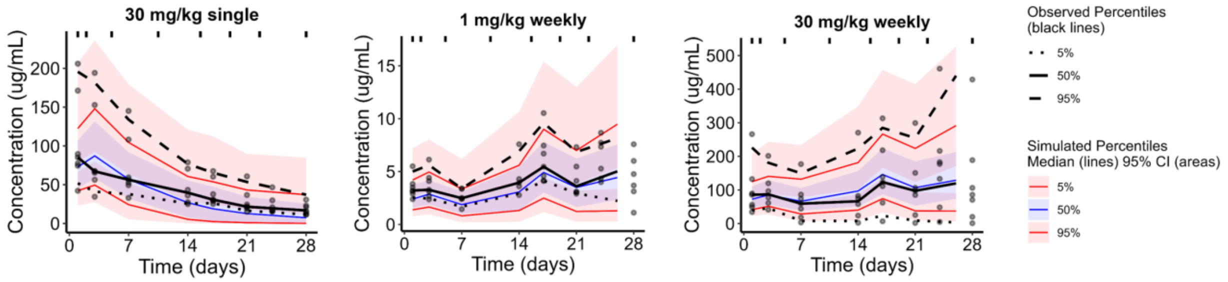


**Figure S4. Visual predictive check stratified by pilot arm** (chAdu 30 mg/kg single dose, 1 mg/kg once weekly, 30 mg/kg once weekly), based on 1000 simulated replicates of the model-building dataset using the final-model fixed and random effects. Black lines and symbols are observed 5th, 50th, and 95th percentiles within prespecified time bins. Shaded regions are 95% prediction intervals around the corresponding simulated percentiles.

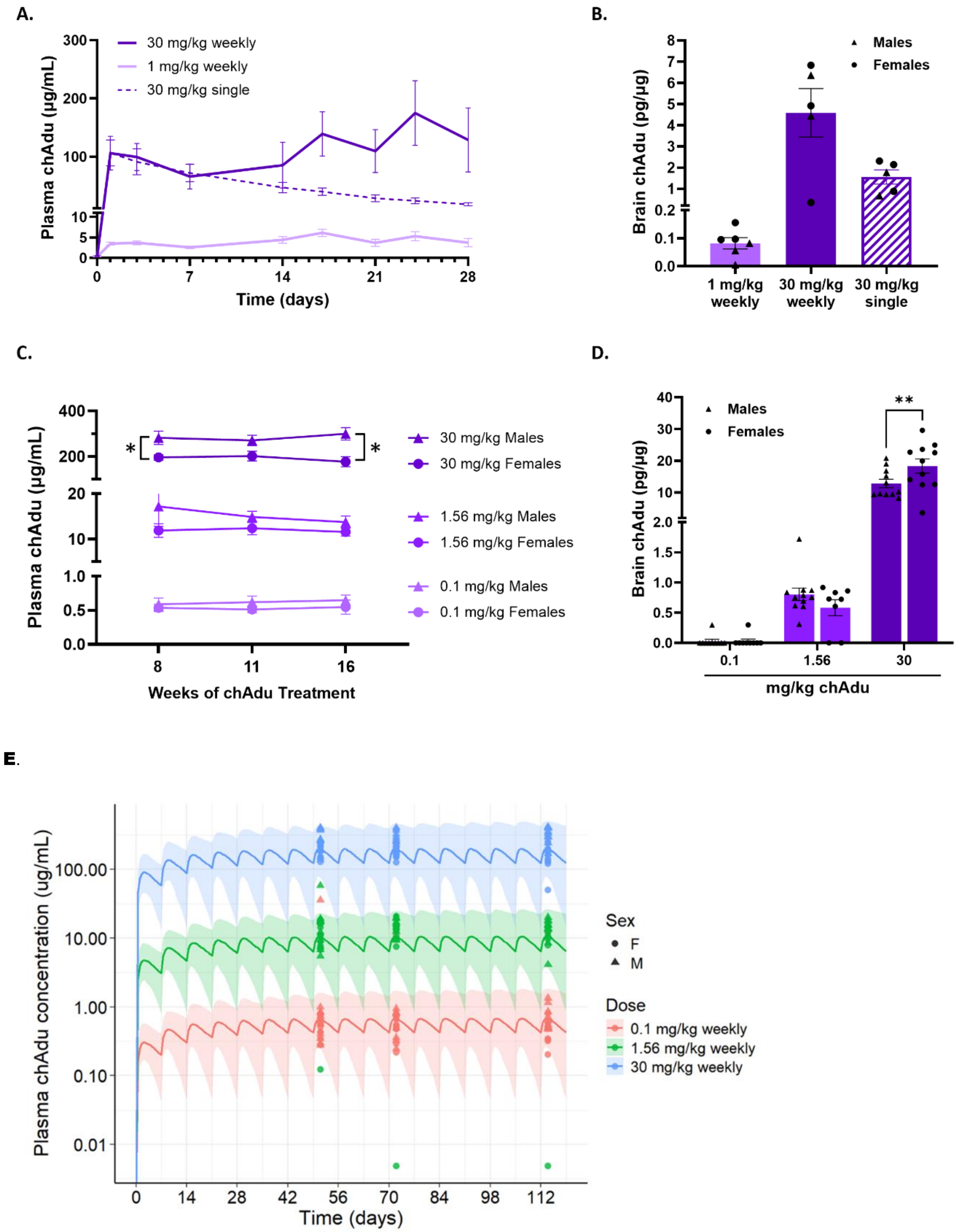


| Regimen | $C_{max,ss}$ (µg/mL) | $C_{trough,ss}$ (µg/mL) | $AUC_{\tau,ss}$ (d. µg/mL) |
|---|---|---|---|
| **0.1 mg/kg weekly** | 0.6 | 0.4 | 3.8 |
| **1.56 mg/kg weekly** | 10.0 | 6.3 | 59.3 |
| **30 mg/kg weekly** | 192.9 | 120.6 | 1140.9 |

**Figure S5. Pharmacokinetic analyses including all mice regardless of ADA status.**

Plasma and brain pharmacokinetics (PK) in pilot and chronic dosing cohorts including all animals regardless of anti-drug antibody (ADA) status. Data are presented as mean ± SEM.

(A) Plasma PK in the pilot cohort. (B) Brain PK in the pilot cohort. (C) Plasma PK in the chronic dosing cohort. (D) Brain PK in the chronic dosing cohort. (E) Model-predicted versus observed steady-state plasma chAdu concentrations during chronic dosing. Simulated plasma concentration-time profiles for once-weekly intraperitoneal chAdu administration at 0.1, 1.56, and 30 mg/kg over 17 weeks. Solid lines represent the population median from 500 simulated subjects using final popPK model parameter estimates with between-subject variability (BSV) on CL/F and V/F; shaded regions represent 90% prediction intervals. Individual observed chronic cohort plasma chAdu concentrations (circles, females; triangles, males) are overlaid at weeks 8, 11, and 17 (48 hours post-dose).

Analyses were performed as described for Figure 1B-E, with animals grouped by assigned treatment irrespective of ADA status. Animals identified as misdosed based on detectable drug levels in terminal plasma or brain in IgG- or saline-treated groups were excluded *a priori* and removed from all analyses.

To evaluate the impact of anti-drug antibodies (ADA) on pharmacokinetic outcomes, PK analyses were repeated including all animals regardless of ADA status (Supplementary Fig. S5). Overall patterns were consistent with those observed in the primary analyses (Fig. 1B-E), indicating that exclusion of ADA+ animals did not materially alter PK interpretations. Subjects with treatment-emergent ADAs (10-19% incidence; see Fig. 1F-G) may contribute to low-exposure observations.

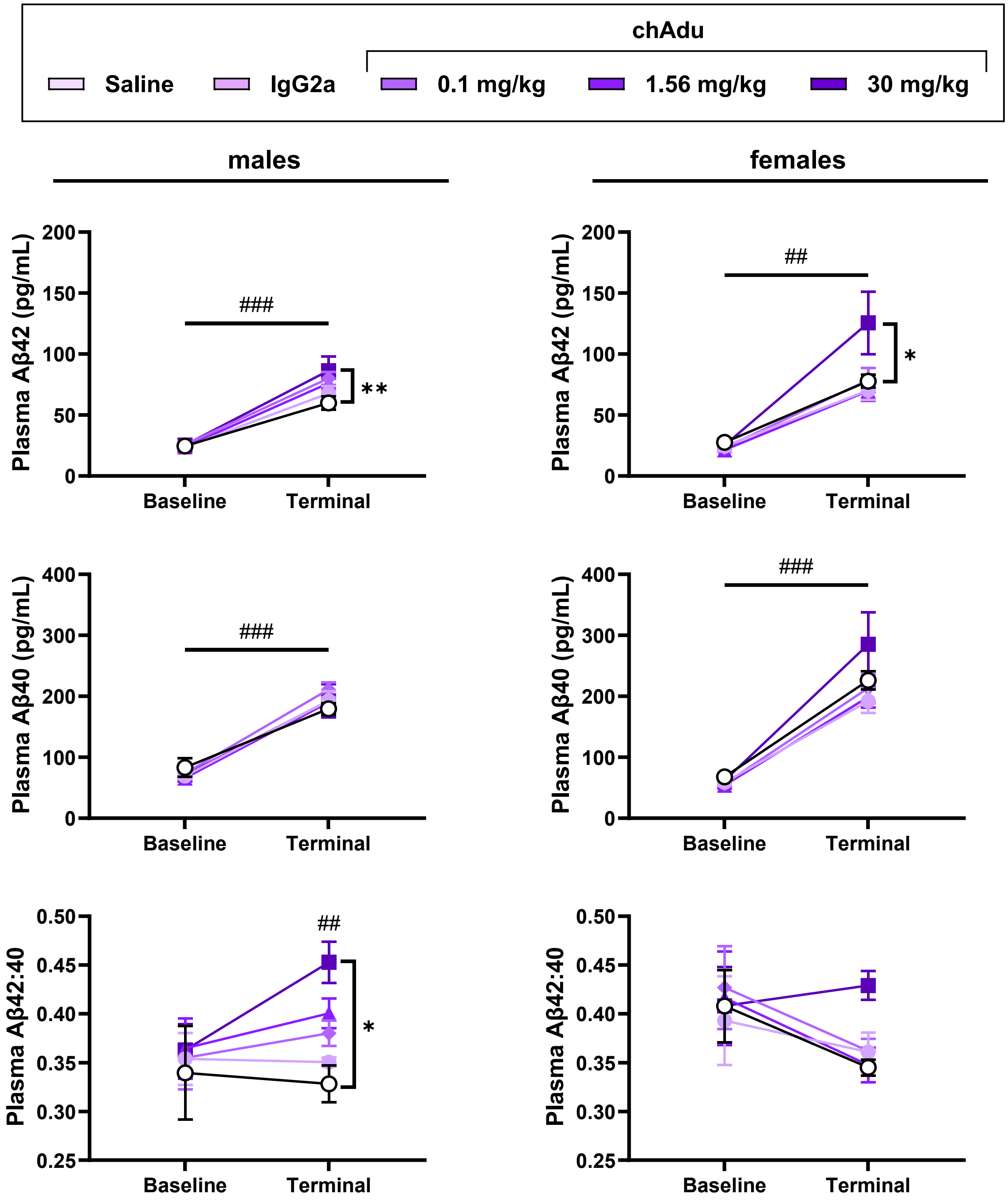


**Figure S6. Plasma pharmacodynamic (PD) outcomes including all mice regardless of ADA status**

Plasma amyloid-β (Aβ) measures in male and female mice including all animals regardless of anti-drug antibody (ADA) status. Data are presented as mean ± SEM.

(A–B) Plasma Aβ42:40 ratios in males (A) and females (B). (C–D) Plasma Aβ42 levels in males (C) and females (D). (E–F) Plasma Aβ40 levels in males (E) and females (F).

Points indicate group mean ± SEM. Data were analyzed within each sex using two-way repeated-measures ANOVA (time × treatment) followed by Tukey's multiple comparisons test. Statistical annotations indicate: * and **, differences between treatment groups at the terminal timepoint; #, significant differences between baseline and terminal within treatment group(s). Where # is shown above a single group, the effect applies only to that group; where shown as a spanning bar, the effect applies across all groups (*$p < 0.05$, **$p < 0.01$; #$p < 0.05$, ##$p < 0.01$, ###$p < 0.001$).

To evaluate the impact of anti-drug antibodies (ADA) on pharmacodynamic outcomes, analyses were repeated including all animals regardless of ADA status (Supplementary Fig. S6). Overall patterns of treatment effects were consistent with those observed in the primary analyses (Fig. 2), indicating that ADA-based exclusions did not substantially alter study conclusions. Inclusion of ADA-positive animals resulted in minor changes in statistical significance in select comparisons.

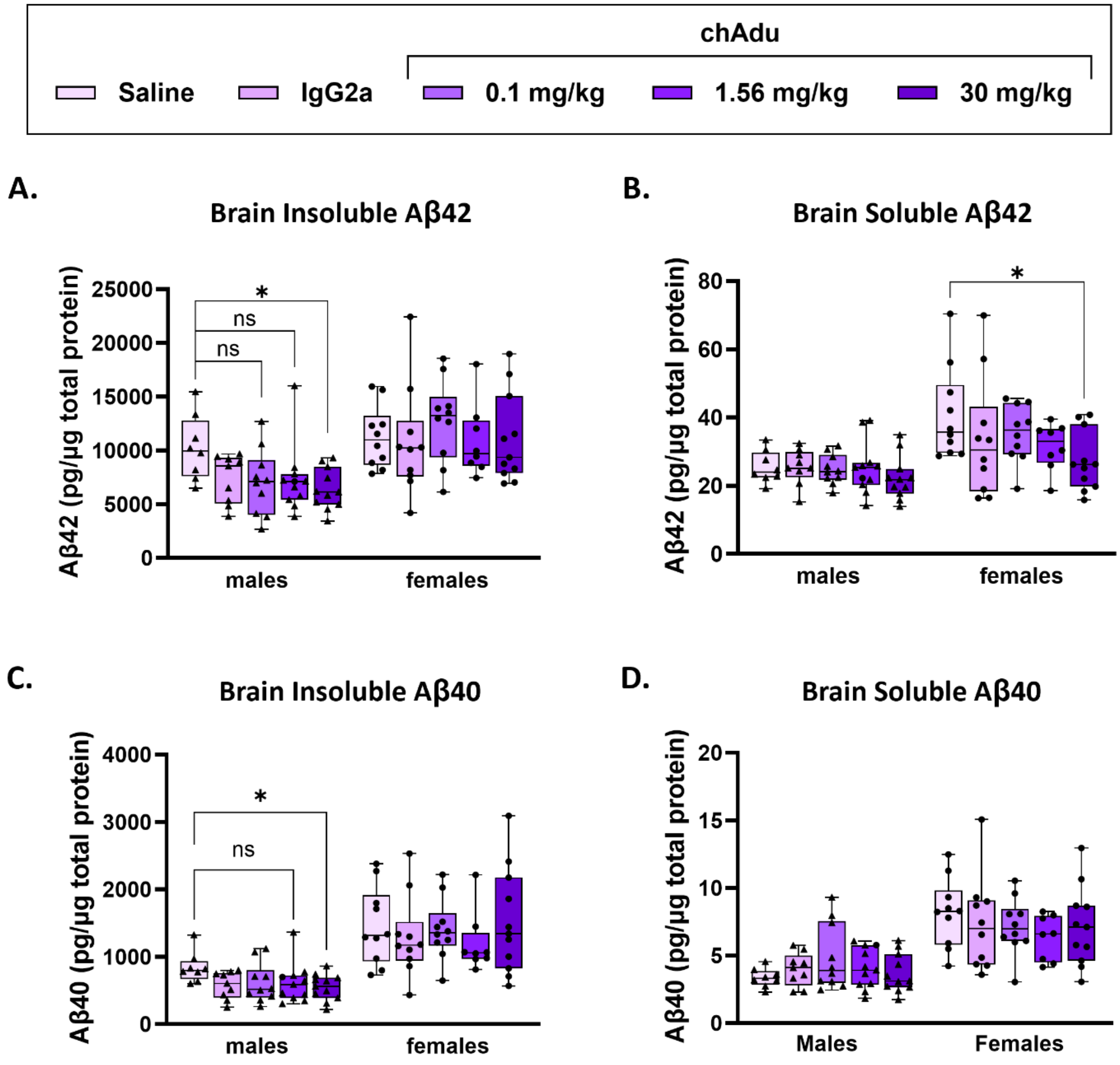


**Figure S7. Brain Aβ species following chAdu treatment including all mice regardless of ADA status**

(A) Insoluble Aβ42 in males and females. (B) Soluble Aβ42 in males and females. (C) Insoluble Aβ40 in males and females. (D) Soluble Aβ40 in males and females.

Data are presented as box-and-whisker plots showing the median and interquartile range; whiskers indicate minimum to maximum values. Individual data points represent single animals (triangles, males; circles, females).Within each sex, data were analyzed by one-way ANOVA followed by Dunnett's multiple comparisons test, with IgG- and chAdu-treated groups compared to saline-treated controls. Statistical annotations indicate significant differences relative to saline-treated animals. Animals were grouped by assigned treatment irrespective of ADA status.

To determine whether inclusion of ADA-positive animals influenced brain Aβ outcomes, analyses were repeated including all animals regardless of ADA status (Supplementary Fig. S7). Overall patterns of treatment effects on soluble and insoluble Aβ species were consistent with those observed in the primary analyses (Fig. 3), with only small changes in statistical outcomes (Supplementary Fig. S7A, C).

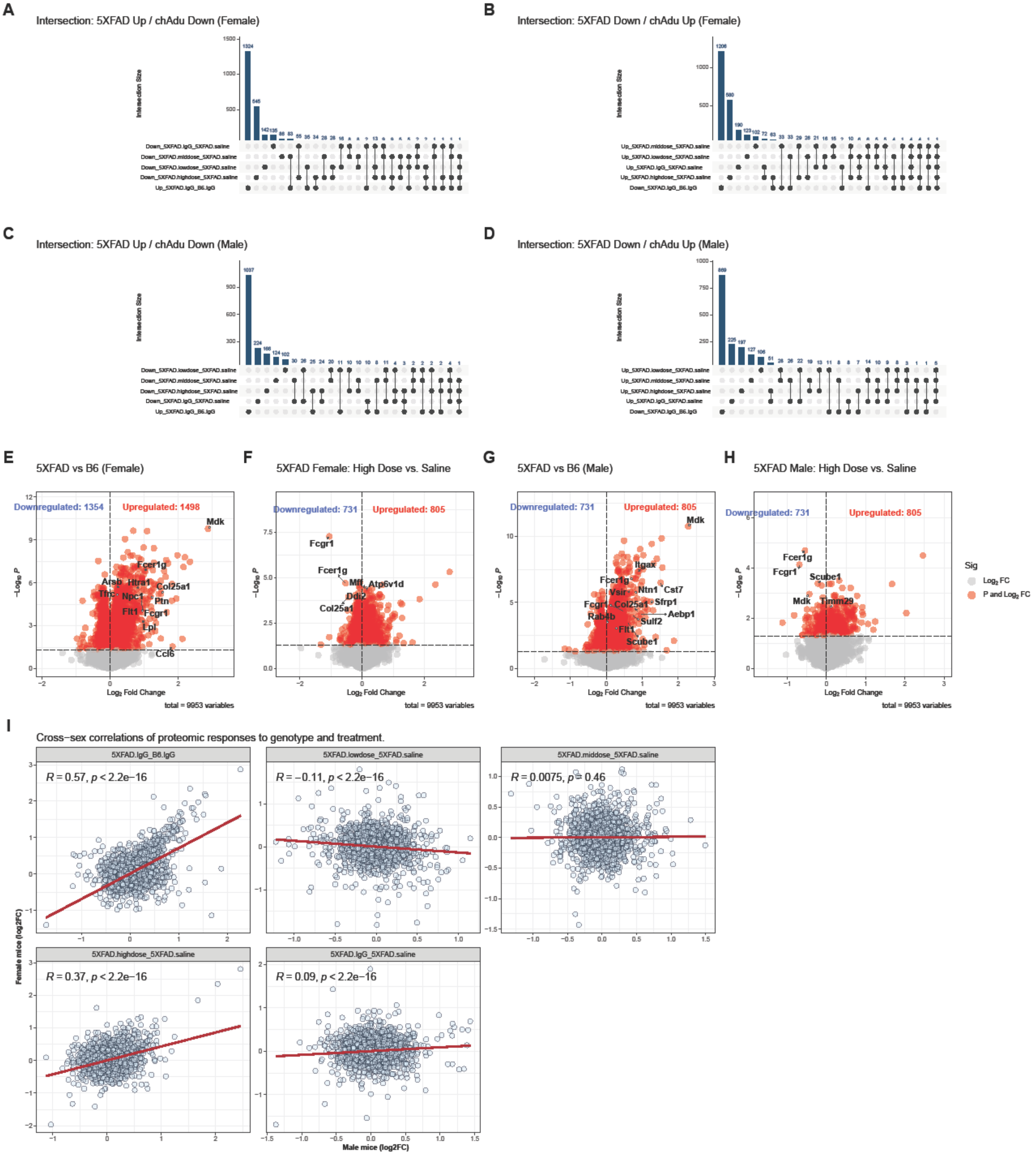
A
Intersection: 5XFAD Up / chAdu Down (Female)
B
Intersection: 5XFAD Down / chAdu Up (Female)
C
Intersection: 5XFAD Up / chAdu Down (Male)
D
Intersection: 5XFAD Down / chAdu Up (Male)
E
5XFAD vs B6 (Female)
Downregulated: 1354
Upregulated: 1498
F
5XFAD Female: High Dose vs. Saline
Downregulated: 731
Upregulated: 805
G
5XFAD vs B6 (Male)
Downregulated: 731
Upregulated: 805
H
5XFAD Male: High Dose vs. Saline
Downregulated: 731
Upregulated: 805
Log2 Fold Change
total = 9953 variables
Sig
Log2 FC
P and Log2 FC
I
Cross−sex correlations of proteomic responses to genotype and treatment.
5XFAD.IgG_B6.IgG
R = 0.57, p < 2.2e−16
5XFAD.lowdose_5XFAD.saline
R = −0.11, p < 2.2e−16
5XFAD.middose_5XFAD.saline
R = 0.0075, p = 0.46
5XFAD.highdose_5XFAD.saline
R = 0.37, p < 2.2e−16
5XFAD.IgG_5XFAD.saline
R = 0.09, p < 2.2e−16
Female mice (log2FC)
Male mice (log2FC)

**Figure S8: Comparative intersection and correlation analysis of sex-specific proteomic responses to chAdu and IgG treatment.**

**(A, C) Intersections of disease-associated upregulation and treatment-mediated downregulation.** UpSet plots illustrating the overlap between proteins significantly upregulated in IgG2a-treated 5XFAD mice relative to IgG2a-treated WT littermates and those downregulated following chAdu treatment in (A) females and (C) males (FDR < 0.05). **(B, D) Intersections of disease-associated downregulation and treatment-mediated upregulation.** UpSet plots depicting the overlap between proteins significantly downregulated in IgG2a-treated 5XFAD mice relative to IgG2a-treated WT littermates and those upregulated following chAdu or IgG2a treatment in (B) females and (D) males (FDR < 0.05). **(E, G) Baseline proteomic shifts in isotype-control cohorts.** Volcano plots depicting differentially expressed proteins (DEPs) in IgG2a-treated 5XFAD mice relative to IgG2a-treated WT littermates in (E) females and (G) males. Red points indicate significantly up- or downregulated proteins (FDR < 0.05); grey points denote proteins meeting the fold-change threshold but not statistical significance. **(F, H) High-dose chAdu-mediated proteomic remodeling.** Volcano plots displaying significantly up- or downregulated proteins (FDR < 0.05) in high-dose chAdu-treated 5XFAD mice compared to saline-treated 5XFAD controls for (F) females and (H) males. Proteins exhibiting significant directional reversal—those upregulated by 5XFAD genotype but downregulated by treatment, or vice versa—are explicitly labeled. Red points indicate significance (*FDR < 0.05 and abs(log2FC)>0*|), while grey points indicate proteins meeting only the fold-change criteria. **(I) Cross-sex correlations of proteomic responses to genotype and treatment.** Scatter plots illustrate the Pearson correlation (*R*) of protein expression changes ($log_2FC$) between female (y-axis) and male (x-axis) 5XFAD mice across different experimental conditions. Each point represents an individual protein, with red regression lines indicating the direction and strength of the linear relationship.
